# Domain-general and language-specific contributions to speech production in a second language: an fMRI study using functional localizers

**DOI:** 10.1101/2023.07.02.547419

**Authors:** Agata Wolna, Jakub Szewczyk, Michele Diaz, Aleksandra Domagalik, Marcin Szwed, Zofia Wodniecka

## Abstract

For bilinguals, speaking in a second language (L2) compared to the native language (L1) is usually more difficult. In this study we asked whether the difficulty in L2 production reflects increased demands imposed on domain-general or core language mechanisms. We compared the brain response to speech production in L1 and L2 within two functionally-defined networks in the brain: the Multiple Demand (MD) network and the language network. We found that speech production in L2 was linked to a widespread increase of brain activity in the domain-general MD network. The language network did not show a similarly robust differences in processing speech in the two languages, however, we found increased response to L2 production in the language-specific portion of the left inferior frontal gyrus (IFG). To further explore our results, we have looked at domain-general and language-specific response within the brain structures postulated to form a Bilingual Language Control (BLC) network. Within this network, we found a robust increase in response to L2 in the domain-general, but also in some language-specific voxels including in the left IFG. Our findings show that L2 production strongly engages domain-general mechanisms, but only affects language sensitive portions of the left IFG. These results put constraints on the current model of bilingual language control by precisely disentangling the domain-general and language-specific contributions to the difficulty in speech production in L2.

## Introduction

Bilinguals often struggle with speaking in their second language (L2) compared to their native language (L1), even when they have a good level of proficiency. This difficulty is reflected in increased naming times and decreased accuracy in tasks requiring participants to name pictures of simple objects in L1 and L2 [1–5]. Similarly, neuroimaging research using *EEG* [6–7] and *fMRI* [8–13] consistently shows that speech production in L2 is linked to increased engagement of neural resources.

In this study, we investigated the neural correlates of L1 and L2 speech production to gain a better understanding of the source of the difficulty accompanying speech production in L2. Specifically, we asked whether production in L2 entails increased engagement of domain-general control resources, the core language processes, or both. Even though brain mechanisms engaged in speech production in L1 and L2 have been studied before, we are still lacking a full understanding of the mechanisms that contribute to speech production in the non-native language. One of the central axes of this debate is whether the increased effort related to speaking in L2 can be traced back to increased engagement of domain-general or language-specific processes. However, addressing this problem is a non-trivial task as the assumptions of the theoretical models of bilingual speech production as well as the tools used to interpret the brain responses to L2 and L1 do not clearly disentangle the language-specific and domain-general mechanisms engaged in speech production. Speech production in both L2 and L1 needs to rely on language-specific processes (conceptual preparation, lexical access, phonological encoding, articulation), however differences in the brain response to naming pictures in L2 and L1 usually have been attributed to increased engagement of cognitive control, not core language mechanisms. This interpretation draws on the assumption of theoretical models of bilingual speech production that speaking in a weaker language requires increased engagement of bilingual language control mechanisms that modulate the activation of the two languages and adjust them to the task and contextual demands at hand [14–16]. The problem is that from this perspective, all differences in neural activity between speech production in L2 and L1 should be interpreted as increased engagement of the control mechanisms. Therefore, this approach can easily overlook differences in language-specific mechanisms supporting the processing of L1 and L2, as the theoretical framework of bilingual language production does not differentiate between the domain-general and language-specific components of the postulated control mechanisms. In the current paper, we explored the brain response to speech production in L1 and L2 using an analytical approach in which functional networks are defined independently of the main task. This approach precisely disentangles the domain-general and language-specific components of bilingual speech production.

### Speech production in L2 as a difficulty for domain-general control mechanisms

Many studies have linked the difficulty related to speech production in L2 to increased engagement of the domain-general control mechanisms [15,17–20]. The cognitive processes orchestrated by the bilingual language control network largely overlap with those attributed to the Multiple Demand (MD) network [21] in both their functional definitions and anatomical localization in the brain. The MD network is a domain-general fronto-parietal network that supports a wide range of executive control mechanisms [22], working memory [23], fluid intelligence and problem solving [24]. The MD network also engages in the execution of complex and non-automated tasks [23,25,26] such that its activation closely reflects the difficulty of a given task [23]. This network is characterized by broad domain-generality [27–28] and can be simultaneously engaged in many aspects of a given task, including sensory (visual, auditory and motor) demands and rule complexity [27]. It has been suggested that the role of the MD network is to integrate information that is relevant to a current cognitive operation [29,30]. Importantly, although the MD network is sensitive to integration across a broad array of tasks, it is not strongly engaged by basic language processes (e.g., sentence comprehension [23,31]) Since L2 is typically less automatized than L1, the sensitivity of the MD network to task automatization, makes it a suitable candidate for supporting the additional demands related to producing speech in L2. This possibility is further corroborated by the topographical overlap between the MD network and structures that form the bilingual language control network.

### Speech production in L2 as a difficulty for core language mechanisms

In addition to increased task demands, the difficulty related to speaking in L2 may reflect a language-processing problem and entail increased demands for language-specific computations in the brain which are supported by the language network: a fronto-temporal neural network specifically tuned to process language [31,32]. The language network supports representational and computational aspects of language processing. It is engaged in language comprehension [32,33] and production [34] across different modalities and tasks. It supports language processing at the sub-lexical [35], lexical, and syntactic levels [36–38], it is also sensitive to the difficulty of language processing [39–40] and responds to differences between comprehension in native and non-native languages in polyglots [41]. What is more, the language network was shown to be sensitive to lexical access demands [34], and lexical access difficulty related to speaking in the L2 compared to the native language [42]. As such, it is plausible to assume that speech processing in L2 may present a language processing difficulty that is handled by the language network.

### What do we know about the neural basis of speech production in L2?

Neuroimaging research has shown that production in L2 compared to L1 engages a wide network of bilateral cortical and subcortical brain structures including the left and right prefrontal cortex, dorsal anterior cingulate cortex and pre-supplementary motor area, left and right inferior parietal lobe, cerebellum, thalamus as well as basal ganglia [15,20]. Abutalebi and Green [15] proposed that these structures form the Bilingual Language Control network that supports a variety of processes necessary for a bilingual speaker to efficiently use two languages. The bilingual language control network has been identified predominantly based on studies using the language switching tasks (LST [8,9,17,43–47]; for meta-analyses see [13,48]). However, these findings may be difficult to interpret as the LST captures brain activations underlying speech production in L1 and L2 in a situation requiring the speakers to switch between languages very frequently (e.g., every few seconds), which is not typically experienced in everyday life. Such frequent switches may engage cognitive control to a much higher degree than language production in more naturalistic situations. Specifically, frequent switching between the two languages may lead to increased brain response that not only reflects the engagement of neural networks supporting the L1 and L2 but also the mechanisms that support execution of difficult, non-automated tasks [25,26]. Finally, picture naming in L1 is usually significantly slowed down in the LST, which is interpreted as a consequence of rebalancing activation levels of L1 and L2 (the *reverse dominance* phenomenon [49]). As such, comparison of brain responses to picture naming in L1 and L2 in the LST may reflect artificial task demands that may even reverse language dominance due to switching between languages, rather than mechanisms that support production in L1 and L2 in a less demanding, single language context.

Then, it is not surprising that studies comparing language production in L1 vs L2 in single-language context show a different pattern of brain activity than those identified using the LST [13]. This discrepancy in results is also consistent with predictions of the adaptive control hypothesis of bilingual language control [18] which assumes that cognitive control resources are flexibly adjusted to the communicative context and therefore they may be differentially engaged in different tasks. For these reasons, it is essential to examine the brain correlates of bilingual speech production in a single language context. So far, several studies have used this approach [9–12,33,44,50–52]. However, their results do not converge on a set of brain regions responding more strongly to speech production in L2 compared to L1 (see **Table 1** and **Figure 1**). While the brain regions identified in these studies mostly pertain to the bilingual language control network [15], there is little overlap in peaks of activation across the studies. What is more, because of a high individual variability in the precise topography of functional networks supporting high-level cognitive processes [53,54], it is difficult to say the activations observed in the previous experiments fall within the language or the MD networks (this is especially problematic for activations falling within brain regions where language and MD networks are adjacent to each other).

**Figure 1.**
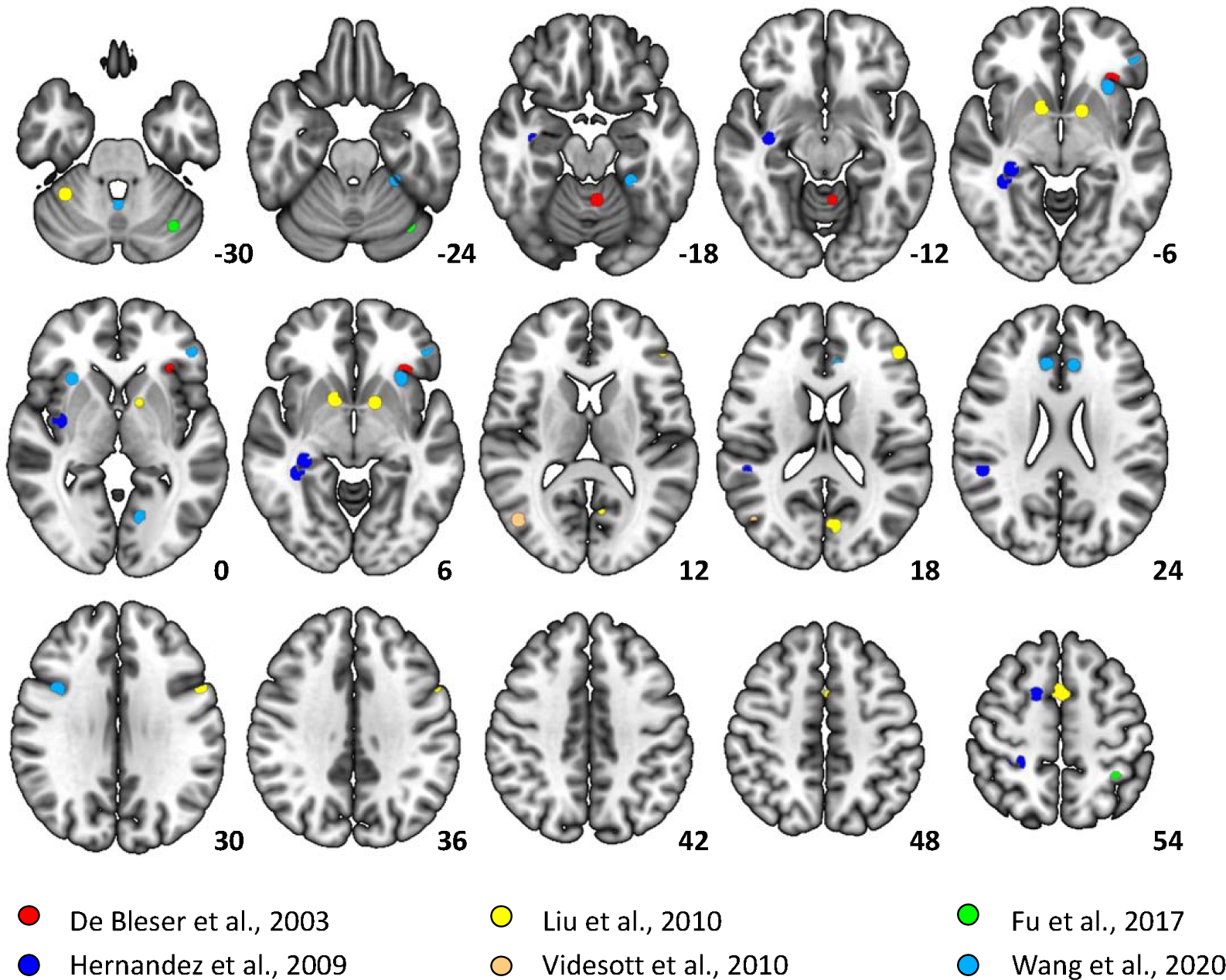
Peak activations corresponding to the L2 > L1 contrast in previous studies looking at differences between languages in single-language context. Data was plotted using a 5mm radius around each of the peaks reported in the papers. Rossi et al., 2018 did not report the exact peak locations for L2 > L1 contrast, for this reason this study is not included in the figure.

**Table 1.**
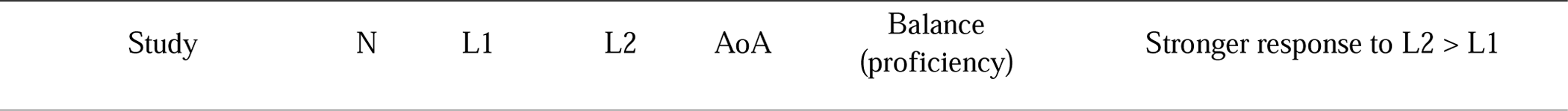
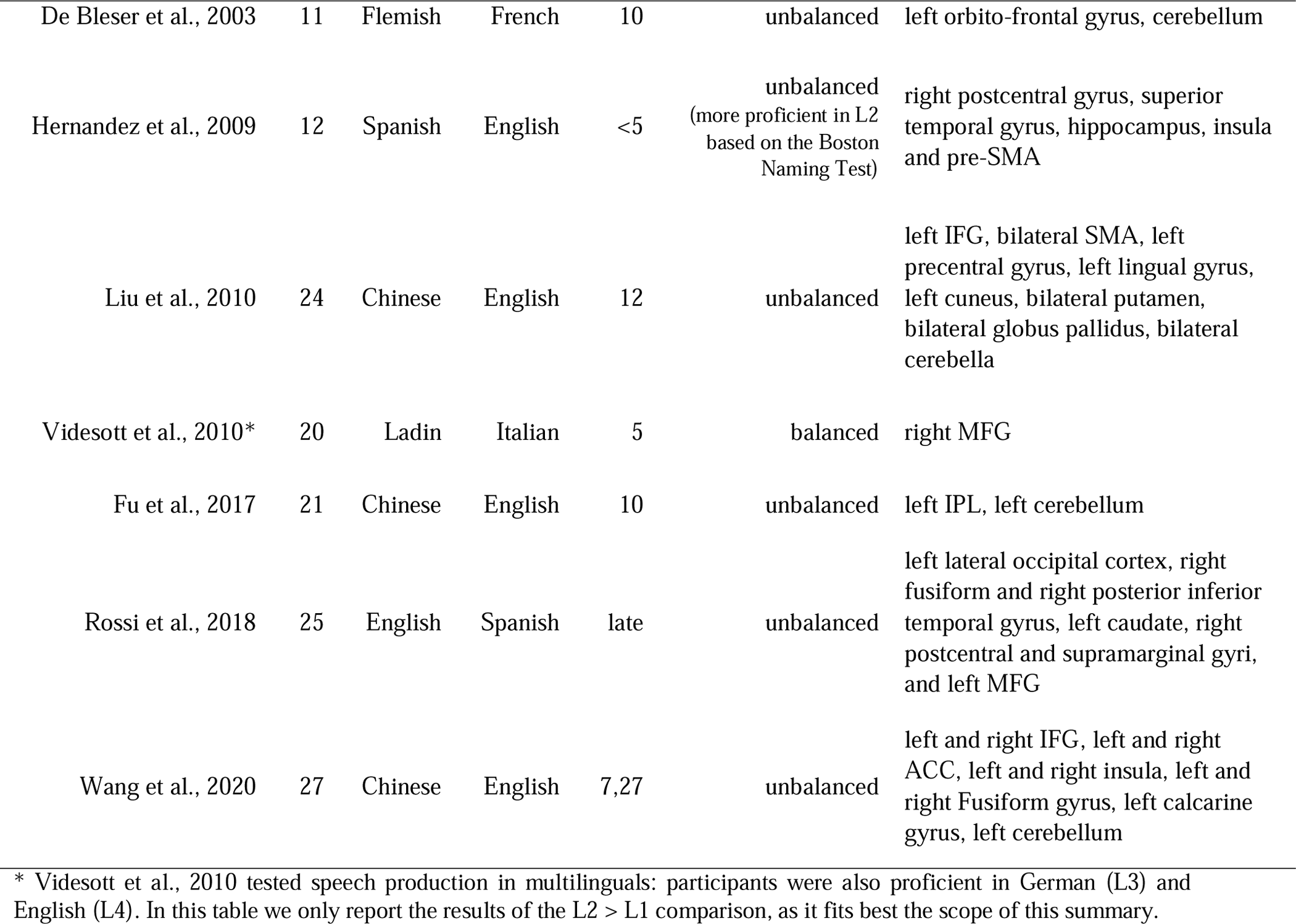
Summary of studies looking at L2 > L1 contrasts in the single language context. The table includes all available studies that tested and reported a direct comparison between L2 and L1. It includes the characteristics of the participants as well as a list of structures that were significantly activated for the L2 > L1 contrast.

### Lateralization of mechanisms supporting speech production in L1 and L2

Another perspective at looking at the brain basis of bilingual speech production is to ask whether within the Language and MD networks L1 and L2 engage similar or different regions. In this context, an interesting yet unanswered question related to the brain basis of speech production in L1 and L2 is the degree of lateralization. It is known that language is lateralized to the left hemisphere in most people, but research suggest that bilinguals may have reduced lateralization, especially if they acquired the L2 early in life [55–58]. However, so far, we know very little about differences in lateralization of L1 and L2 processing. Here we ask two questions related to lateralization of bilingual speech production. First, whether within the language network L1 and L2 show a similar degree of lateralization and second, whether any differences in lateralization of L1 and L2 processing reflect differences in lateralization of language-specific processes or mechanisms supporting the domain-general control.

### Current study

In the current study we asked whether the brain’s solution to processing L2 rely on domain-general control processes or on language specific mechanisms. More specifically, we explored the following questions: Is speech production in L2 linked to increased engagement of domain-general control regions? Is speech production in L2 linked to increased engagement of language-processing regions? Is there a difference in lateralization supporting L1 and L2 production? Additionally, to link our results to the neurocognitive model of bilingual language control [15], we explored brain activations during L1 and L2 speech production within the language-and domain-general selective areas forming the bilingual language control network. To address these questions, we used a precision fMRI approach based on functional localization [32]. To evaluate the brain response to speech production in L2 and L1 within the Language and MD networks, we created a set of subject-specific functional ROIs within each network and estimated the response within these fROIs to the critical conditions: speech production in L2 and L1. Using a similar approach, we created language and domain-general specific fROIs within each brain structure of the bilingual language control network (selected based on [15]; for details see Methods) and estimated responses to speech production in the L2 and L1 within each of the fROIs. A key advantage of this approach is that it precisely identifies functional networks for each subject separately which allows for better modelling of the individual differences in cortical organization of the functional networks in the brain. It also provides a straightforward functional interpretation of the observed brain activity without referring to the function assigned to a particular brain region based on its anatomical label (*reverse inference* problem [59]). Finally, using functional localizers increases the power of statistical tests as the comparisons between L1 and L2 are computed in a narrow set of ROIs instead of thousands of voxels in the entire brain. The increase in statistical power may be particularly important because some of the discrepancies in previous studies’ results may be due to low statistical power.

## Results

### Behavioural results

The analysis of naming latencies revealed that participants were significantly faster in L1 (mean = 1073ms, SD = 233ms) compared to L2 (mean = 1191ms, SD = 264ms). We also found a significant effect of the *Session* showing that participants were overall slower in the 2^nd^ session (mean = 1136ms, SD = 246ms) than in the 1^st^ session (mean = 1095ms, SD = 253ms). Importantly, the effect of *Session* did not interact with the effect of *Language*. The analysis of accuracy yielded very similar results: participants were more accurate in their L1 (mean = 95,1%, SD = 21,5%) than L2 (mean = 66,3% SD = 47,3%), they were also less accurate in the 2^nd^ session (mean = 77,4% SD = 41,8%) than in the 1^st^ session (mean = 83,9%, SD = 36,7%) and there was no interaction between the two effects. The results of the statistical analyses on the behavioural data are presented in **Table 2**.

**Table 2.**
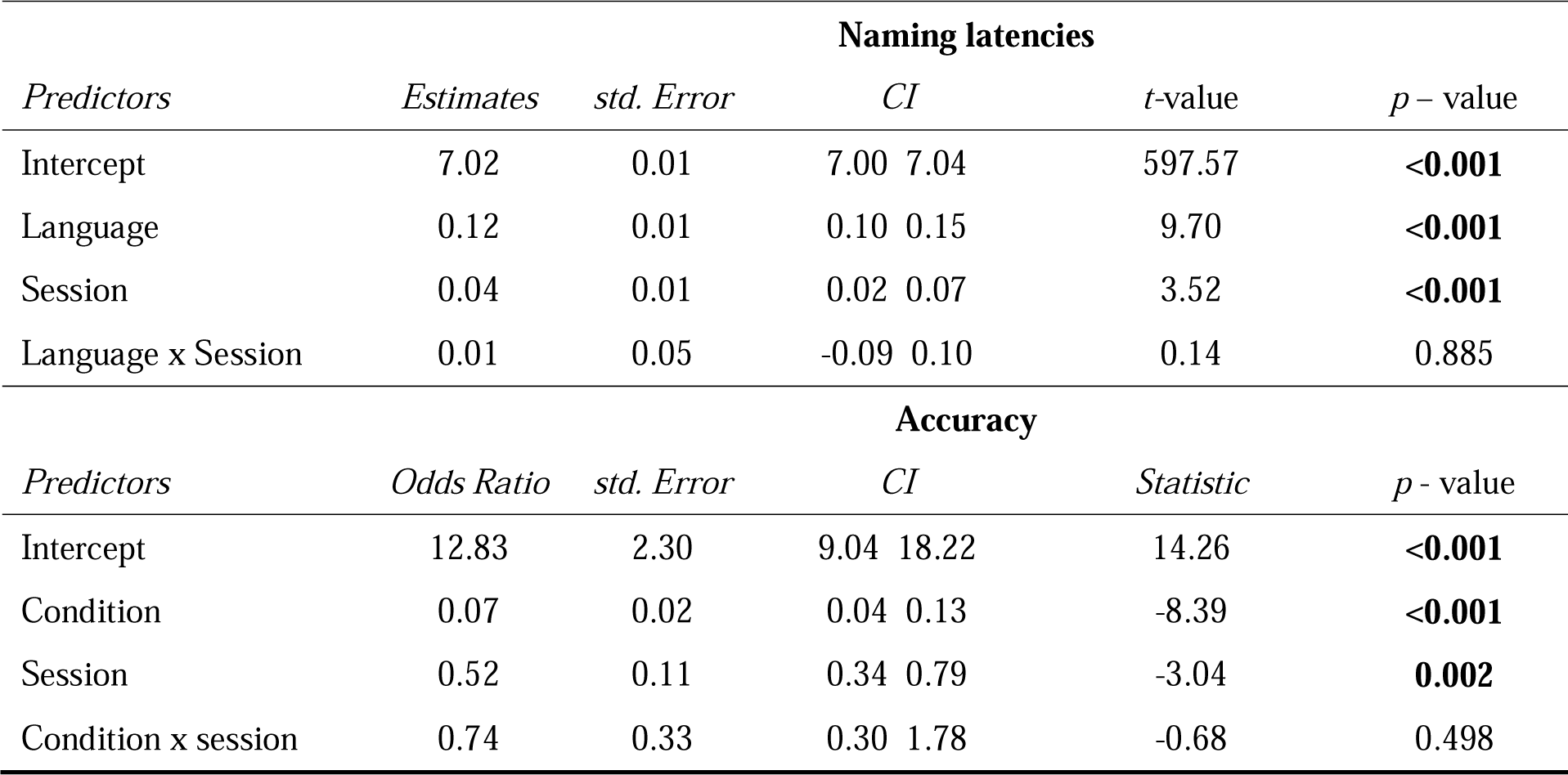
Behavioural performance in Picture Naming task. The table presents results of linear mixed models predicting naming latencies and accuracy of naming pictures.

### Neuroimaging results: localizer approach

In the language network we did not observe a significant effect of *Language* or *Hemisphere*, however, we found a significant interaction between *Language* and *Hemisphere*. Pairwise comparisons revealed that the brain response to speaking in L1 was stronger compared to L2 in the right hemisphere (b = 0.09, *t* = 2.11, *p* = .038) but not different from L2 in the left hemisphere (b = −0.04, *t* = −0.94, *p* = .350). In the MD network, we found a significant effect of *Language* showing stronger responses to L2 than L1 production and *Hemisphere* showing stronger responses in the left than right hemisphere ROIs. Interestingly, we also found an interaction between *Language* and *Hemisphere*. Pairwise comparisons revealed that the response to L2 was stronger than to L1 in the left hemisphere (b = −0.187, *t* = −5.138, *p* < .001) but there were no differences between languages the right hemisphere (b = −0.04, *t* = −0.994, *p* = .325). The results of the localizer analyses are presented in **Table 3 and Figure 2A & 2C.** To further explore the results, we have also analysed the effect of language (L2 vs. L1) in each fROI separately within both language and MD networks. Within the language network we found increased response to L2 compared to L1 in left IFG (pars opercularis). Within the MD network, significant increase in activation in response to L2 compared to L1 was found in all left hemisphere fROIs (save for the left midParietal fROI) and in the right insula and SMA. The results of the by-fROI analyses are presented in **Tables 4 and5** and **Figure 2B & 2D**.

**Figure 2.**
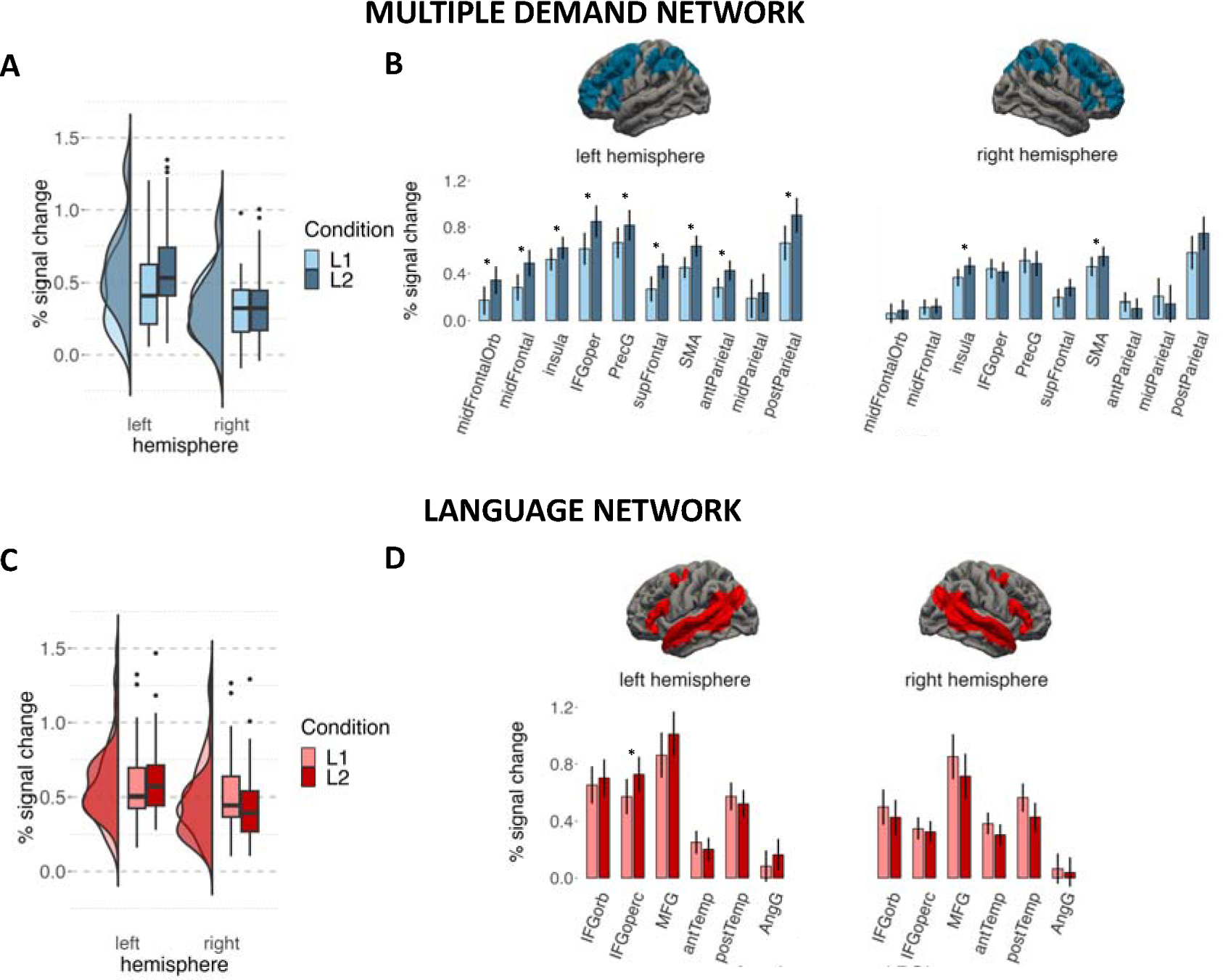
Brain response to speech production in L1 and L2 within the MD and language networks. The plot presents estimates of % signal change predicted by the statistical models. Panels A) and C) present data for estimates to L1 and L2 within the MD and language networks averaged by hemisphere. Individual data points in these panels correspond to predicted responses of individual subjects; and panels B) and D) represent means for naming in L1 and L2 for each ROI. Whiskers in these panels represent Confidence Intervals for post-hoc pairwise comparisons, and significant differences between conditions for a given ROI are marked with asterisks.

**Table 3.**
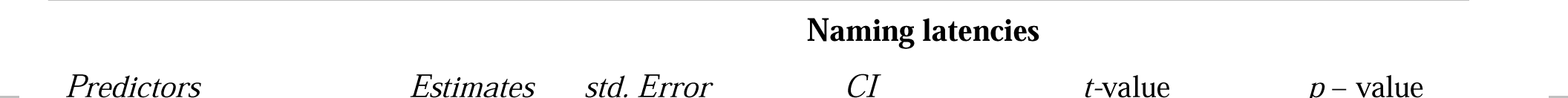
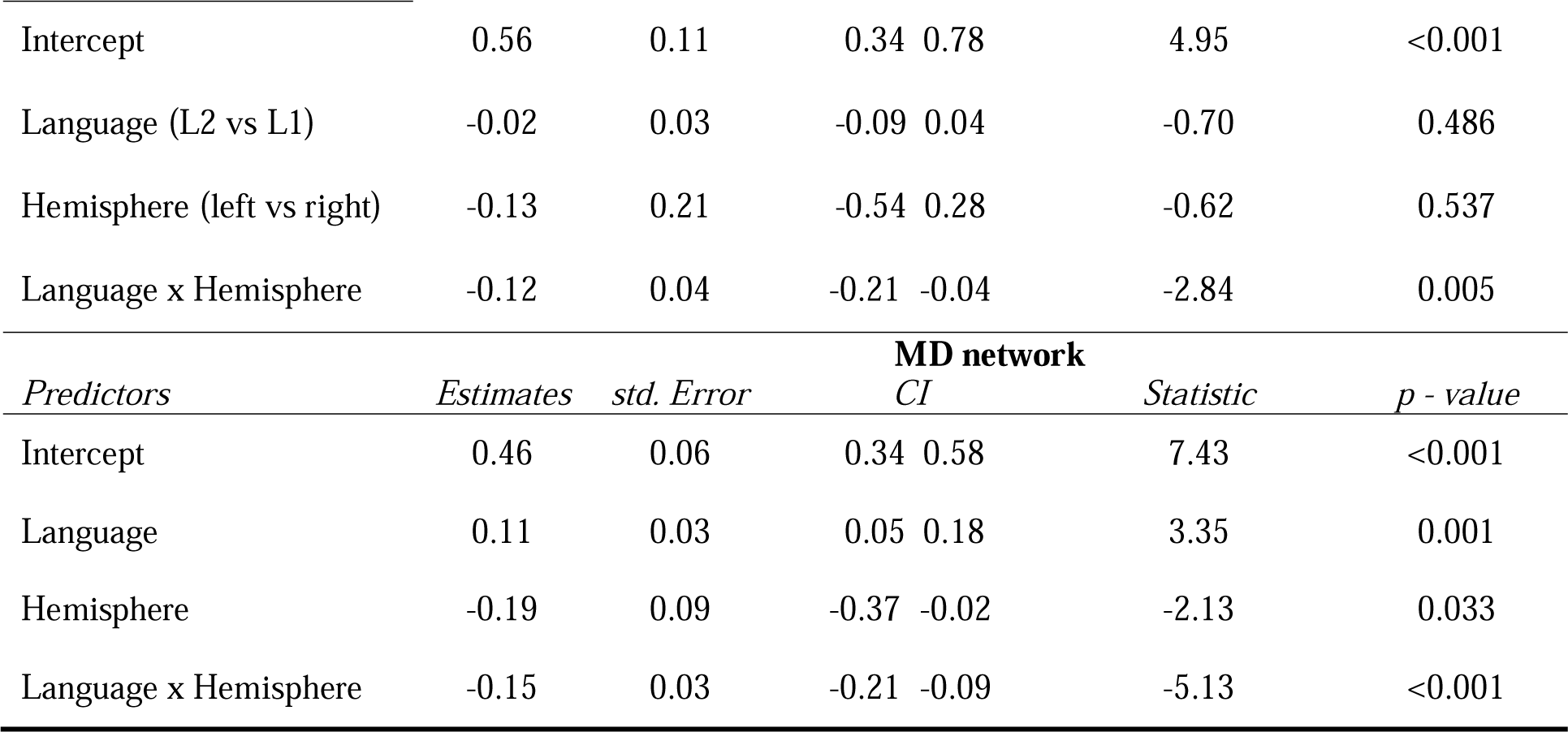
Estimated brain response in Language and MD Network. The table presents results of linear mixed models predicting the % BOLD signal change in the two networks.

**Table 4.**
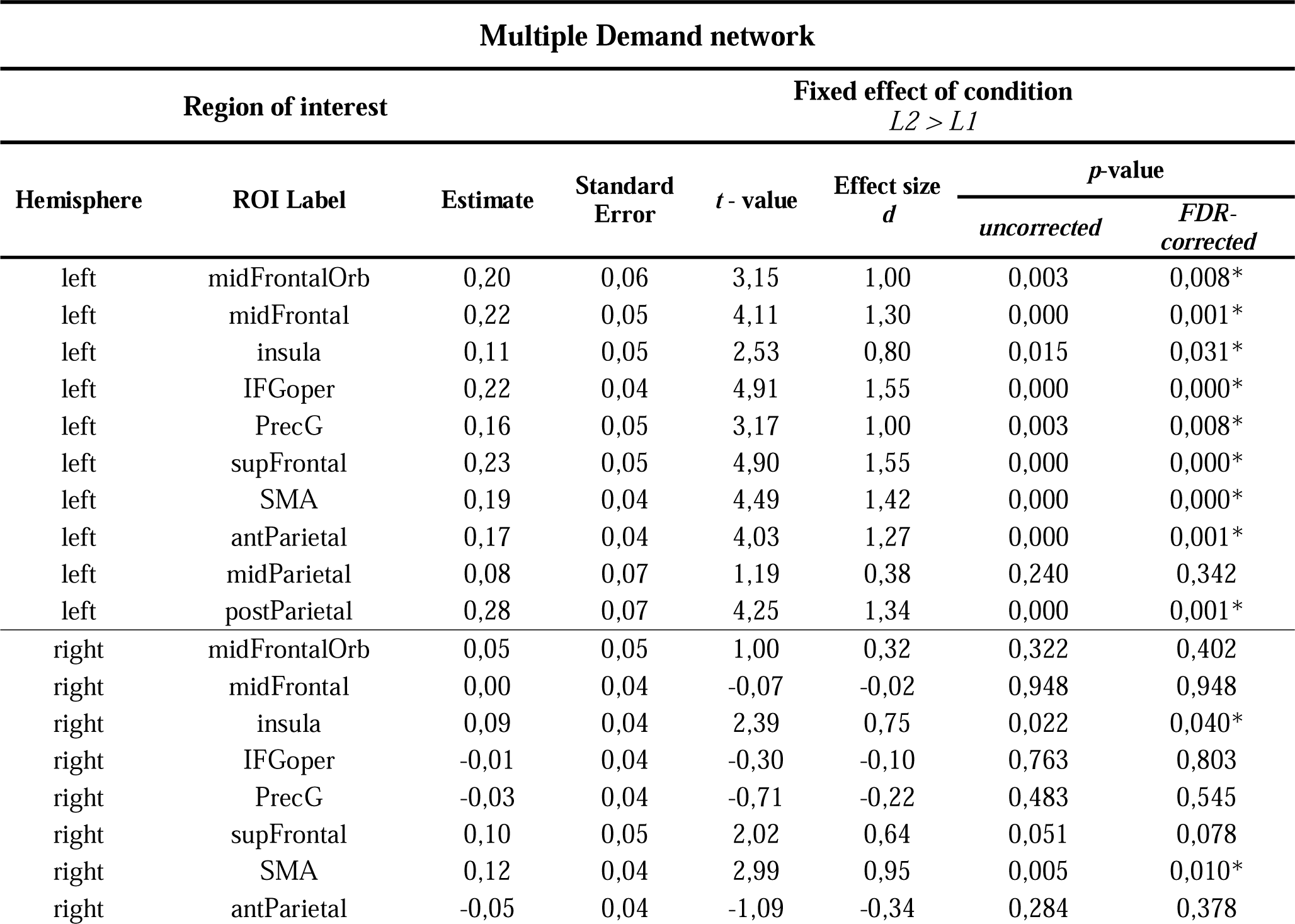
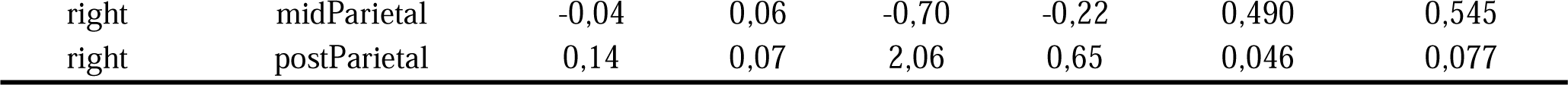
Results of linear mixed-effect models fitted for each functional ROI within the MD network. Presented estimates correspond to the effect of the language (L2 vs L1). Short labels correspond to labels used in Figure 2 in the manuscript. Anatomical labels were derived from Harvard-Oxford Cortical, Harvard-Oxford Subcortical, or Cerebral probabilistic atlases (from FSL) and they correspond to one or two labels with the highest probabilistic overlap with each functional ROI.

**Table 5.**
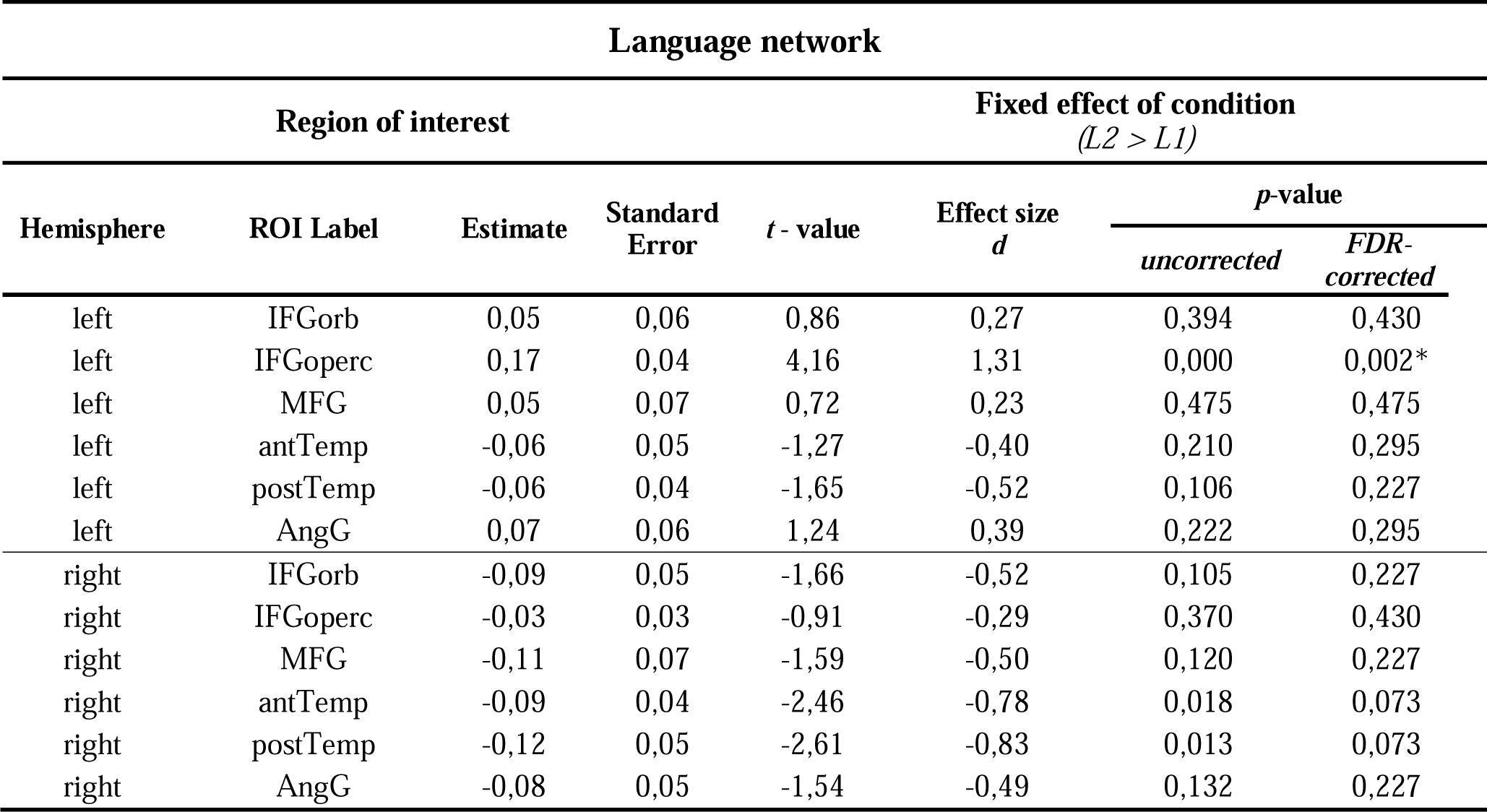
Results of linear mixed-effect models fitted for each functional ROI within the language network. Presented estimates correspond to the effect of the language (L2 vs L1). Short labels correspond to labels used in Figure 2 in the manuscript. Anatomical labels were derived from Harvard-Oxford Cortical, Harvard-Oxford Subcortical, or Cerebral probabilistic atlases (from FSL) and they correspond to one or two labels with the highest probabilistic overlap with each functional ROI.

To provide a more direct link between brain activation and the difficulty of speech production, we have analysed the relationship between brain response in the Language and MD networks with mean reaction times (RTs) in a picture naming task in L1 and L2, separately. In the language network we found no effect of the RTs (b = −0.06, t = −0.33, p = .746), the hemisphere (b = −0.13, t = −0.63, p = .542) or the interaction between RTs and hemisphere (b = −0.29, t = −1.47, p = .144). In the MD network, however, we found a main effect of RTs (b = 0.76, t = 3.49, p = .002), hemisphere (b = −0.20, t = −2.19, p = .041) and the interaction between the RTs and hemisphere (b = −0.61, t = −4.256, p < .001) reflecting that slower RTs were related to larger increase in brain activation in the left than in the right hemisphere (for detailed results, including the by-ROI analyses see the Supplementary Materials). The results of this analysis are presented in Figure 3.

**Figure 3.**
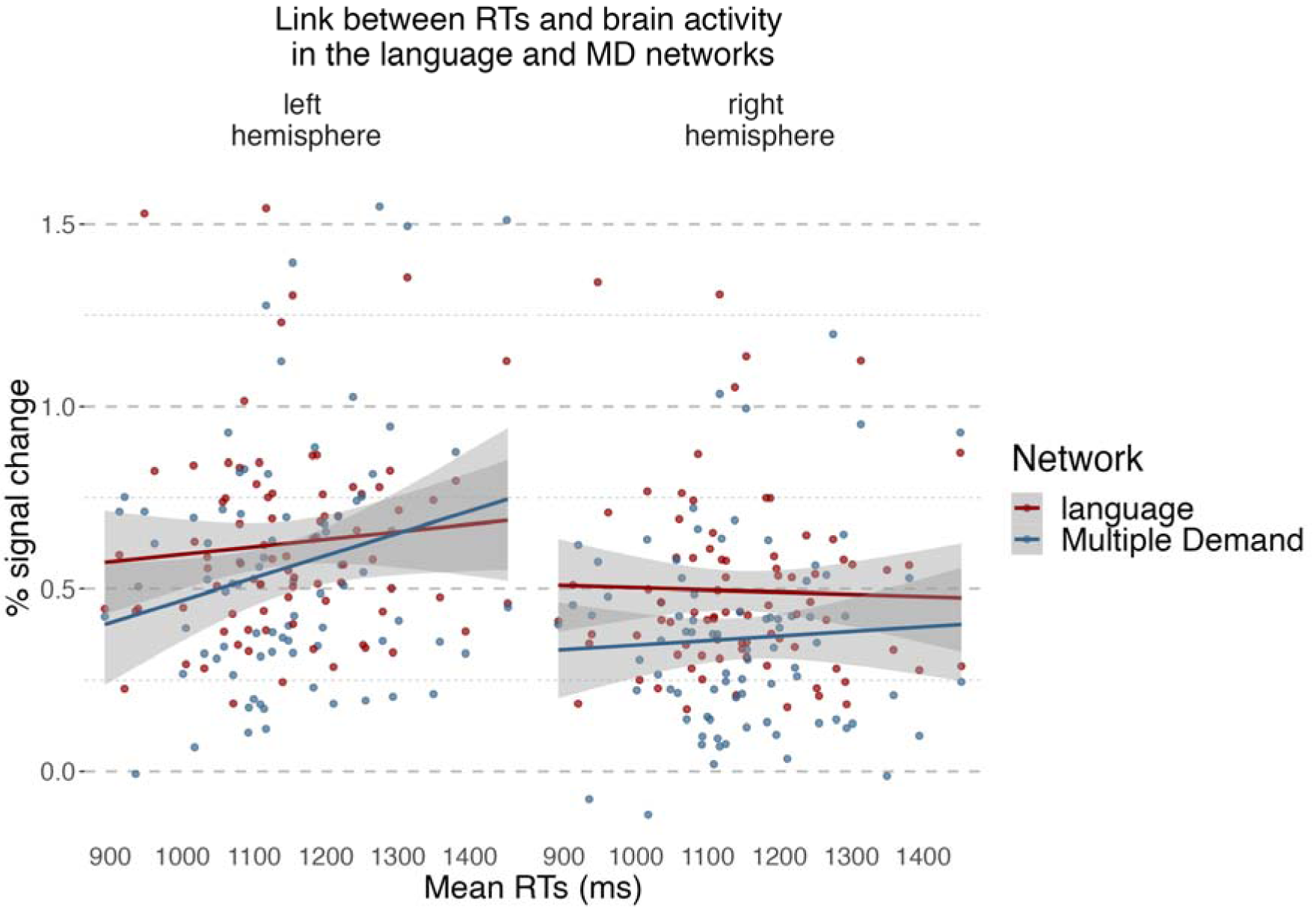
Relationship between naming latencies corresponding to naming pictures and brain response in the language and MD networks. The plot presents a relationship between naming latencies in L2 and L1 (averaged for each language and each participant) and % signal change for each condition in the language and MD networks.

Finally, we have also explored whether the differences in brain activation between speech production in L2 and L1 are modulated by the individual differences, including the language experience, of the participants. In both the language and the MD networks we found that the brain activity increased with the years of education (Language: b = 0.05, *t* = 2.69, *=* 0.011, MD: b = 0.06, *t* = 3.45, *=* 0.002). Additionally, in the MD network, we found an interaction between language, hemisphere, and proficiency level in L2 (b = 0.79, *t* = 2.01, *p* = 0.037) showing that in the left hemisphere differences in brain response to L2 vs L1 decrease with increasing proficiency in L2 (details of these analyses are reported in the Supplementary Materials in Tables S2-4 and Figure S2).

### Neuroimaging results: bilingual language control network

The first step in the analysis of language-specific and domain-general contributions to the bilingual language control network, was to assess the functional selectivity of language and MD fROIs within the bilingual language control network ROIs. To this aim, we evaluated responses to the localizer contrasts within both language and MD fROIs. Similar to the localizer-based analyses within the language and MD networks, the fROIs were created by taking 10% of most responsive voxels for a given localizer contrast in each of the anatomically defined ROI forming the bilingual language control network (henceforth: BLC network ROIs). Subsequently, the selectiveness of these fROIs to language and domain-general computations was evaluated on an independent portion of the data by comparing their response to the language and MD localizer contrasts. Among the MD fROIs, we observed a robust response to the MD localizer contrasts (hard > easy visual working memory task) in all fROIs except for the left putamen (*p* = .061). The MD fROIs were also found to be highly selective to the domain-general task (i.e., we did not observe significant responses to the language localizer contrast in any of them). For the language fROIs, we observed significant response to the language localizer contrast (listening to intact > degraded speech) in the left angular gyrus, left IFG (pars triangularis and opercularis), left MFG, left pre-SMA and right IFG (pars triangularis and opercularis). What is more, the left MFG fROI was sensitive to not only the language localizer contrast but also to the MD contrast, which means it is not specifically-selective for language processing. Full results of these analyses are presented in the Supplementary Materials in the Table S6. We also found that the overlap between language- and domain-general selective fROIs (in the BLC network ROIs for which both types of fROIs were successfully identified) was very small – the average overlap at the voxel level was not larger than 10% (see Figure 3).

In the next step, the language and MD fROIs that showed a significant response to their localizer contrasts were used to evaluate the brain response to L2 and L1 speech production. Response to L2 was significantly stronger than to L1 in 4 language fROIs and 7 MD fROIs. In the language-selective fROIs we found significant effects in the left IFG (pars triangularis and opercularis), left MFG and left pre-SMA. In the MD-selective fROIs we found increased response to L2 compared to L1 in the left ACC and left pre-SMA, left IFG (pars opercularis and triangularis), left MFG, and left caudate. The results of this analysis are presented in Table 6 and Figure 4.

**Figure 4.**
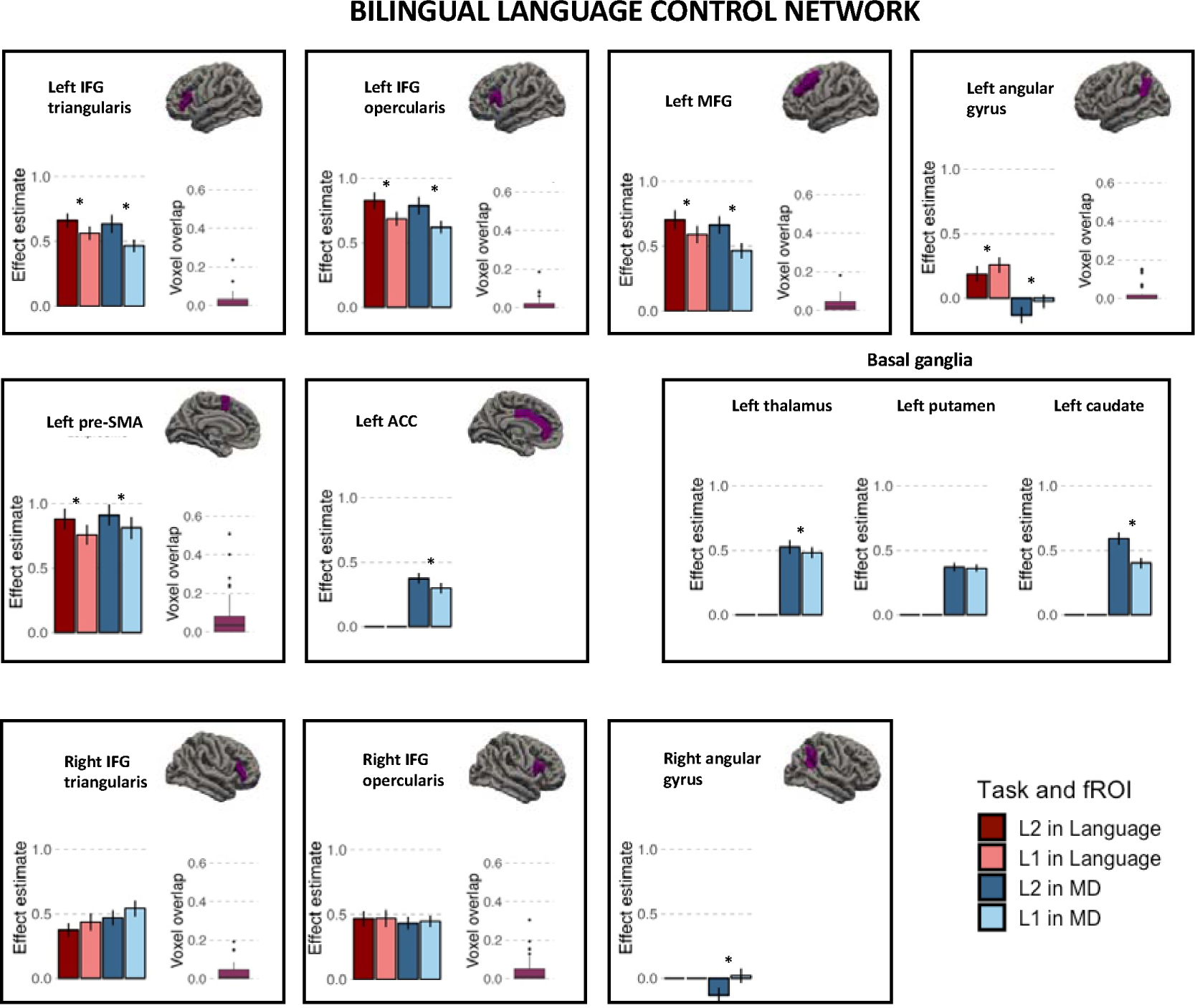
Brain response to speech production in L1 and L2 within the bilingual language control network ROIs. Each panel presents the estimates for speech production in L2 and L1 in the language-specific fROIs (red) and MD-specific fROIs (blue) within a given BLC ROI. For ROIs in which we identified both language- and MD-selective parcels, we also show the % of the voxels overlapping between the two fROIs.

**Table 6.**
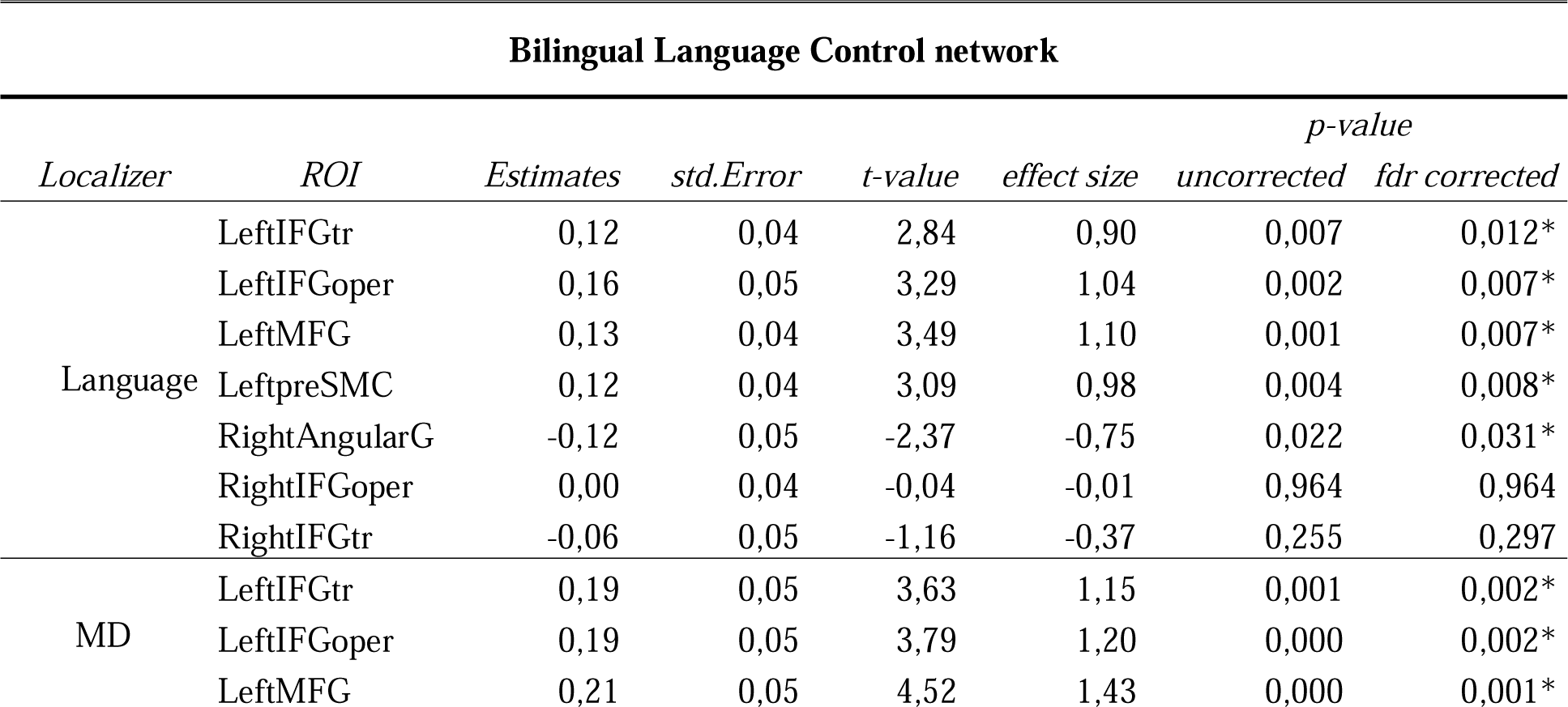
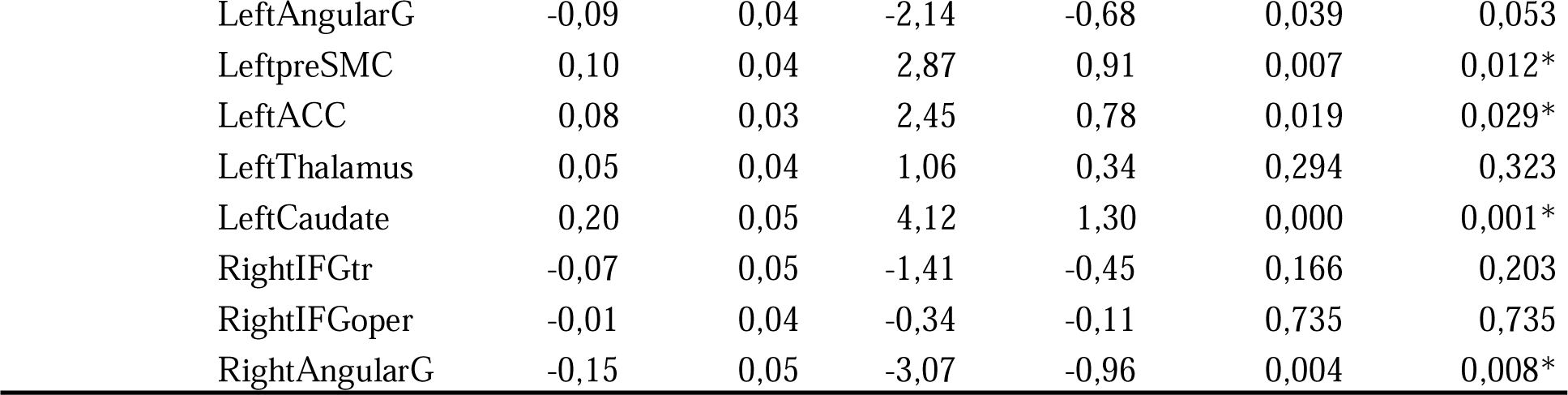
Difference between estimated brain response to speech production in L2 and L1 within language and MD fROIs in the Bilingual Language Control network ROIs. The table presents results of linear mixed models predicting the % BOLD signal change corresponding to L2 > L1 difference in each fROI.

| Bilingual Language Control network |  |  |  |  |  |  |  |
| --- | --- | --- | --- | --- | --- | --- | --- |
| Localizer | ROI | Estimates | std.Error | t-value | effect size | p-value |  |
|  |  |  |  |  |  | uncorrected | fdr corrected |
| Language | LeftIFGtr | 0,12 | 0,04 | 2,84 | 0,90 | 0,007 | 0,012* |
|  | LeftIFGoper | 0,16 | 0,05 | 3,29 | 1,04 | 0,002 | 0,007* |
|  | LeftMFG | 0,13 | 0,04 | 3,49 | 1,10 | 0,001 | 0,007* |
|  | LeftpreSMC | 0,12 | 0,04 | 3,09 | 0,98 | 0,004 | 0,008* |
|  | RightAngularG | -0,12 | 0,05 | -2,37 | -0,75 | 0,022 | 0,031* |
|  | RightIFGoper | 0,00 | 0,04 | -0,04 | -0,01 | 0,964 | 0,964 |
|  | RightIFGtr | -0,06 | 0,05 | -1,16 | -0,37 | 0,255 | 0,297 |
| MD | LeftIFGtr | 0,19 | 0,05 | 3,63 | 1,15 | 0,001 | 0,002* |
|  | LeftIFGoper | 0,19 | 0,05 | 3,79 | 1,20 | 0,000 | 0,002* |
|  | LeftMFG | 0,21 | 0,05 | 4,52 | 1,43 | 0,000 | 0,001* |

|  |  |  |  |  |  |  |
| --- | --- | --- | --- | --- | --- | --- |
| LeftAngularG | -0,09 | 0,04 | -2,14 | -0,68 | 0,039 | 0,053 |
| LeftpreSMC | 0,10 | 0,04 | 2,87 | 0,91 | 0,007 | 0,012* |
| LeftACC | 0,08 | 0,03 | 2,45 | 0,78 | 0,019 | 0,029* |
| LeftThalamus | 0,05 | 0,04 | 1,06 | 0,34 | 0,294 | 0,323 |
| LeftCaudate | 0,20 | 0,05 | 4,12 | 1,30 | 0,000 | 0,001* |
| RightIFGtr | -0,07 | 0,05 | -1,41 | -0,45 | 0,166 | 0,203 |
| RightIFGoper | -0,01 | 0,04 | -0,34 | -0,11 | 0,735 | 0,735 |
| RightAngularG | -0,15 | 0,05 | -3,07 | -0,96 | 0,004 | 0,008* |

## Discussion

In this paper we explored the brain basis of the difficulty accompanying speech production in L2 and L1. Although some previous research has attempted to explore this issue before, there are several strengths and novelties of our contribution. First, we used newer analytical tools – the precision fMRI method based on functional localizers [32,60]. More specifically, we tested the overlap of brain responses to speech production in L2 and L1 with two functional networks in the brain: the language network, which supports core language processes [32] and the MD network, which supports a wide range of domain-general control mechanisms [21,28,32]. The second novelty and strength of our contribution is that, unlike most of the previous studies - we have focused on speech production in the single language context – i.e., we used a task that does not require participants to frequently switch between languages. As such, our findings allow making conclusions about mechanisms of pure L2 vs. L1 production, not confounded with the additional language switching requirement. What is more, we have also analysed how the behavioural performance (naming latencies) and individual differences in language experience of the participants modulated the brain response in the language and MD networks. Finally, our study is very well-powered, based on a sample considerably larger (n=41) than any of the previous experiments looking at the same research question.

Using functional localizers, we have shown that difficulty in speech production in L2 is linked to robust activation in areas delineated by the MD localizer, suggesting that L2 leads to a greater recruitment of domain-general resources compared to L1. The language network did not show similarly widespread and consistently stronger responses to L2, except for the left inferior frontal gyrus (IFG) fROI that responded more to speech production in L2 than in L1.

The increased engagement of the MD network in speech production in L2 shows that bilingual language control is achieved by domain-general mechanisms. As indicated earlier, many of the previous studies evaluating and corroborating the assumption of domain-generality of the bilingual language control used the language switching task, which requires frequent switching between languages. Here, we show that in a single language context, production in L2 also draws on domain-general resources. Specifically, compared to L1, speech production in L2 was linked to widespread engagement of the left MD network. This effect appears to be linked to the difficulty that bilingual speakers experience when using their L2, as implied by analyses showing that the increased activity of within the MD network accompanies slower naming. Furthermore, differences in activations in response to L1 and L2 within the left-hemisphere fROIs of the MD network were modulated by proficiency: higher proficiency was associated with overall lower neural responses and smaller differences between languages. The widespread activation within the MD network in response to L2 production suggests that the difficulty in non-native speech production is not driven by an isolated control mechanism. It is not clear, however, *which* exact domain-general mechanisms are more engaged in the L2 production compared to L1.

Within the core language network, speech production in L2 was not linked to a consistent, network-wide increase in activation compared to production in L1. This result is in line with findings showing substantial overlap between brain representations of L1 and L2 [61–65]. It suggests that language-specific processes are largely shared between languages and the most fundamental difference between bilingual and monolingual language processing is not the “language processing” itself. Importantly, we found no differences in how L1 and L2 speech production engaged the left temporal cortices, which have been linked to lexical access [66]. A difficulty at this stage of production has oftentimes been proposed as a locus of differences between L1 and L2 production [8,10,67]. Further support for the notion that it is not the lexical access difficulty that drives the difficulty in speech production in L2, comes from the analyses looking at the modulation of brain response to L2 and L1 by individual differences in language experience. As the lexical access difficulty is modulated by factors such as proficiency, AoA, and patterns of language use, we should see the effects of (at least some of) these variables on brain response to L2 and L1 within the language system. At the same time, even though the language network did not differ in its overall response to speech production in both languages, we found increased response to L2 compared to L1 in the language-specific left IFG fROI. Similar to the results observed in the MD network, brain activations in the language left IFG fROI were also shown to increase with increasing naming latencies. As such, even though speech production in L2 is not linked to a network-wide increase in the engagement of language-specific mechanisms, it locally engages additional resources in the left IFG. Together, our results suggest that difficulty in producing words in L2 is not necessarily related to lexical access, although it may still be possible that this conclusion is limited to a sample of unbalanced but proficient bilinguals and that less proficient speakers, who only start to master a given language, would show differences between L1 and L2 also in the language network.

The left IFG has long been considered one of the key brain regions supporting language processing (in particular, the articulation, cf. Broca’s area). Activity in this structure has been linked to post-lexical compositional processes, such as phonetic encoding, syllabification, or even later processes related to articulatory planning [66,68–70]. In the light of debate on the role of the left IFG in the language and domain-general processes as well as its contribution to controlling the two languages of a bilingual, it is particularly interesting that the increased activation in response to L2 compared to L1 was found in both language-specific and domain-general fROIs within the posterior part of left IFG (pars opercularis). This specific area is hypothesised to play a role of an interface between language representations and articulatory codes executed by the motor cortices [71]. At the same time the left IFG has been suggested to play a crucial role in general selection among the competing representations [72,73]. While this hypothesis was first put forward based on studies focusing on language processing, the engagement of the left IFG in selection of representations has been shown to extend to non-linguistic domains [74].

Importantly, the left IFG, has been shown to be a heterogeneous structure, encompassing specialised language-specific and domain-general subregions [75,76]. Our data confirm this dissociation: language-specific and domain-general fROIs showed a minimal overlap at a subject-level even if the two functional subsections were identified in the same anatomical masks in the BLC network analysis (see voxel overlap plots for the left IFG triangularis and opercularis on Figure 3). On the one hand, our results may indicate that the domain-general and language-specific subsections of the left IFG closely cooperate to select appropriate words for production. On the other hand, however, the language and MD networks were shown to support very different types of processes [76], so it is plausible that the increase in the domain-general and language-specific sub-sections of the left IFG in response to production in L2 reflects two different types of mechanisms. Given the functional characterization of the left IFG discussed above, we hypothesize that the observed increase in activation of the language-specific sub-section of the left IFG in response to speech production in L2 compared to L1 reflects increased difficulty in phonological encoding and articulatory processing [1,4].

The scope of the analyses based on the functional localizers is limited to the activations within the language and MD systems which might leave out important components of the neural system engaged in bilingual language control. To account for this possibility and to directly link our results to the most influential neurocognitive model of bilingual language control (BLC)[15], we ran an additional analysis within the anatomically-defined ROIs postulated to form the BLC network [15]. As the primary aim of our study is to disentangle language-specific and domain-general components of the mechanisms supporting speech production in L2 and L1, in the BLC analysis we capitalized on our functional localizer approach. This allowed us to (1) identify language-specific and domain-general fROIs within the anatomically-defined BLC regions, and (2) test the response of the language-specific and domain-general fROIs to speech production in L1 and L2. We found domain-general selective fROIs in almost all nodes of the BLC network (except for the left putamen), but not all of them responded to the L2 > L1 contrast: significant effects were found only in the left IFG, left MFG, left pre-SMA/ACC, left thalamus, and left caudate. In the remaining fROIs we either observed a de-activation in response to L2 (left and right angular gyrus) or no difference between languages (right IFG triangularis and opercularis). Language-selective fROIs were only found some of the BLC nodes, including bilateral IFG (triangularis and opercularis), left MFG, left angular gyrus and left pre-SMA. Within these language-selective fROIs, increased response to L2 > L1 was found in the left IFG, left MFG, and left pre-SMA. A reversed effect, L1 > L2, was observed in the left angular gyrus. In sum, the structures identified in our analyses largely overlap with those identified in the neurocognitive model of bilingual language control, but they also highlight a few differences. We believe that they can be easily explained by the fact that the BLC model was largely based on studies using the language switching paradigm, whereas in our study, participants named pictures in a single-language context. This allows us to separate structures differentially involved in L1 and L2 production, from structures specific for execution of a task. For instance, in our data we did not observe any differences between brain activations in response to L2 and L1 in the right IFG. Since the right IFG is implicated in cue detection and inhibition of non-target responses [15,18] and inhibition is postulated to be one of the key mechanisms of bilingual language control [14,15,18], the absence of differences in this structure in our study suggests that during production in a single language context, inhibition is not necessarily engaged to the same degree as in contexts requiring frequent switches between languages. At the same time, it has been claimed that bilingual language control in speech production is achieved by a re-configuration of the domain-general network: it draws on the same structures as the ones engaged in domain-general tasks, however, it adjusts the connections and engagement of the particular nodes to the task at hand [20]. Our data partially confirm this claim: we did find increased activation to L2 compared to L1 in most of the domain-general fROIs. However, we have also identified language-specific fROIs within the BLC network that responded more strongly to L2 compared to L1. These results imply that the regions within the BLC network serve dual role, encompassing both language-specific and domain-general processes which cooperate to select appropriate words for production. With methods relying on group averaging, this spatially fine-grained distinction into the two networks may be easily overlooked.

Our results also show that compared to L1 production, L2 production recruits the left-hemisphere to a greater extent in both networks. In the language network, compared to L1, speech production in L2 engaged more resources in the left hemisphere but fewer resources in the right hemisphere. While the left hemisphere is crucial for formulating propositional speech, the right hemisphere supplements it by controlling prosodic and non-propositional components of language [77]. The right hemisphere is also more engaged in automated speech (e.g., reciting months [78]). As such, the stronger engagement of the right hemisphere in the native than non-native language production may be driven by higher automatization of L1 compared to L2, especially in bilinguals who learned the L2 later in life. In the MD network, when speaking in L2 compared to L1 we found stronger activity in the left hemisphere, with no such differences in the right hemisphere. Interestingly, the differences in lateralization were much more pronounced in the MD network than in the language network. This result indicates that the stronger lateralization of L2, does not necessarily reflect the lateralization of language processes themselves but rather the lateralization of domain-general control mechanisms supporting speech production in L2.

To further extend the results of our study, future research should compare the response to L2 and L1 production of the language-specific regions by defining them separately for L1 and L2 at a single-subject level (i.e., by using a localizer task in L1 and L2). The language network constitutes a functionally-integrated system with a similar topography regardless of the language used to identify it [41]. However, using a task in L2 and L1 separately to identify the language-responsive voxels on single-subject level might reveal some more fine-grained differences between processing of the two languages within the language system.

To conclude, our study provides a characterization of brain mechanisms supporting speech production in L2 and L1. We used a methodological approach that allowed us to precisely disentangle the contribution of domain-general and language-specific mechanisms underlying the process of bilingual speech production. We found that the difficulty in speech production in L2, compared to L1, was linked to increased engagement of a wide range of domain-general mechanisms in the MD network. Importantly, the absence of additional activation of the right IFG suggests that if each language is used separately (i.e., without the immediate need to switch them), language inhibition may not be required. The language network did not show similarly widespread and consistently stronger responses to production in L2; however, speaking in L2 compared to L1 was linked to increased engagement of language-specific resources in the left IFG which may reflect increased difficulty in phonological encoding and articulatory processing. Furthermore, we have shown that the structures postulated to form a bilingual language control network encompass both language-specific and domain-general sub-sections which cooperate to enable successful word selection in bilingual speech production. Finally, we found that in both networks the L2 engaged resources lateralized to the left hemisphere, which suggests that the greater lateralization of language processing in bilinguals may in fact reflect lateralization of cognitive control rather than core language mechanisms.

## Method

### Participants

This study reports the results of data collected from the same group of subjects as the one reported Wolna et al [79]. Forty-two Polish-English late bilinguals took part in the study. One participant was excluded due to the excessive head motion (>2mm) in the main fMRI task resulting in a final sample of forty-one participants (31 females, 10 males, mean age = 23.29, SD = 3.24). All participants were Polish native speakers who learned English as their second language. They declared intermediate or advanced proficiency in English (B2 or higher) and obtain at least 20/25 points in the General English Test by Cambridge Assessment (mean score = 21,32, SD = 1,94). Their proficiency in English was also assessed again using LexTale [80] (mean score = 71,58%, SD = 9,91%). Detailed information on the participants’ proficiency, age of acquisition and daily use of the languages (based on a language questionnaire) can be found in Table 7.

**Table 7.**
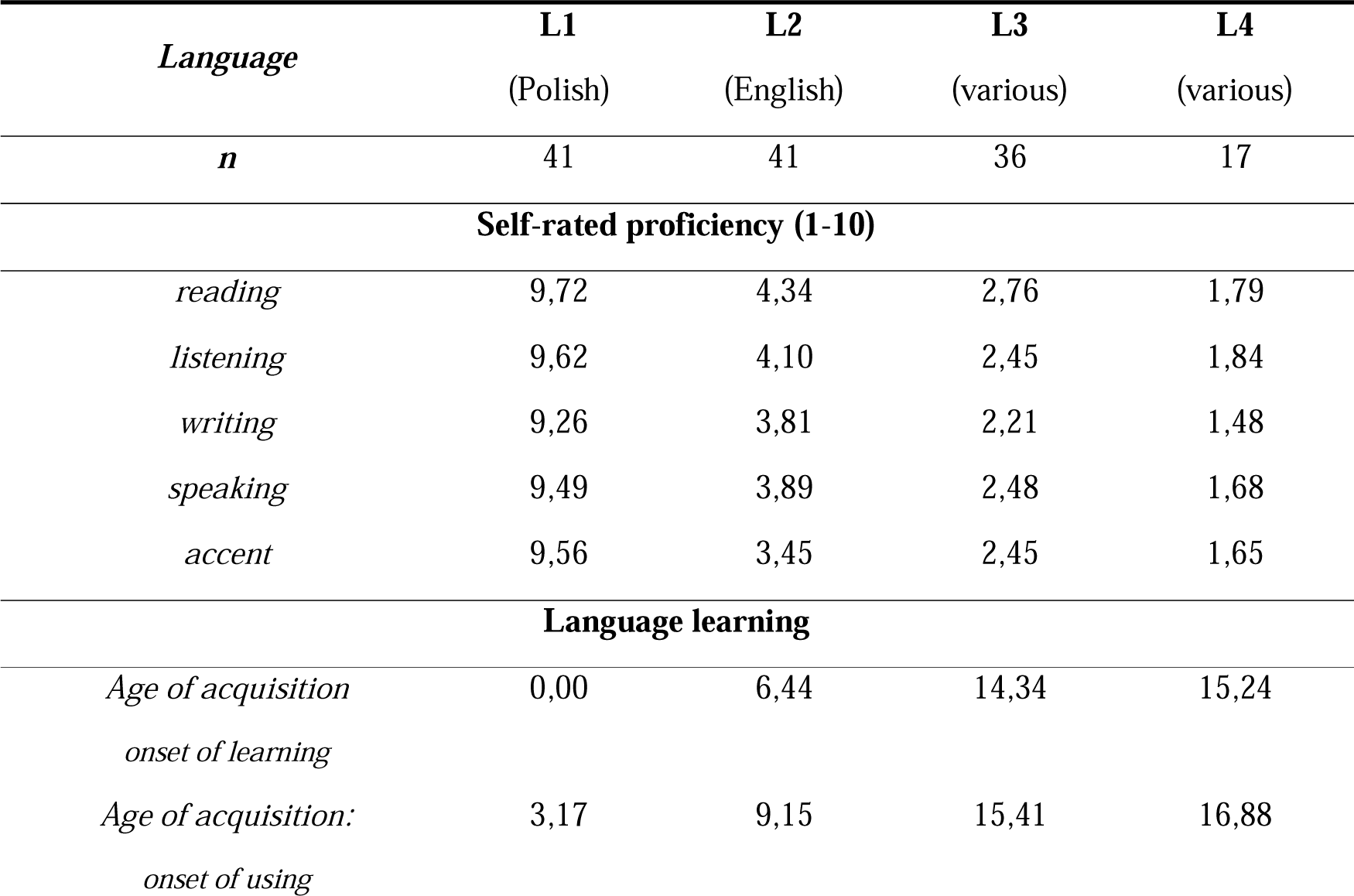
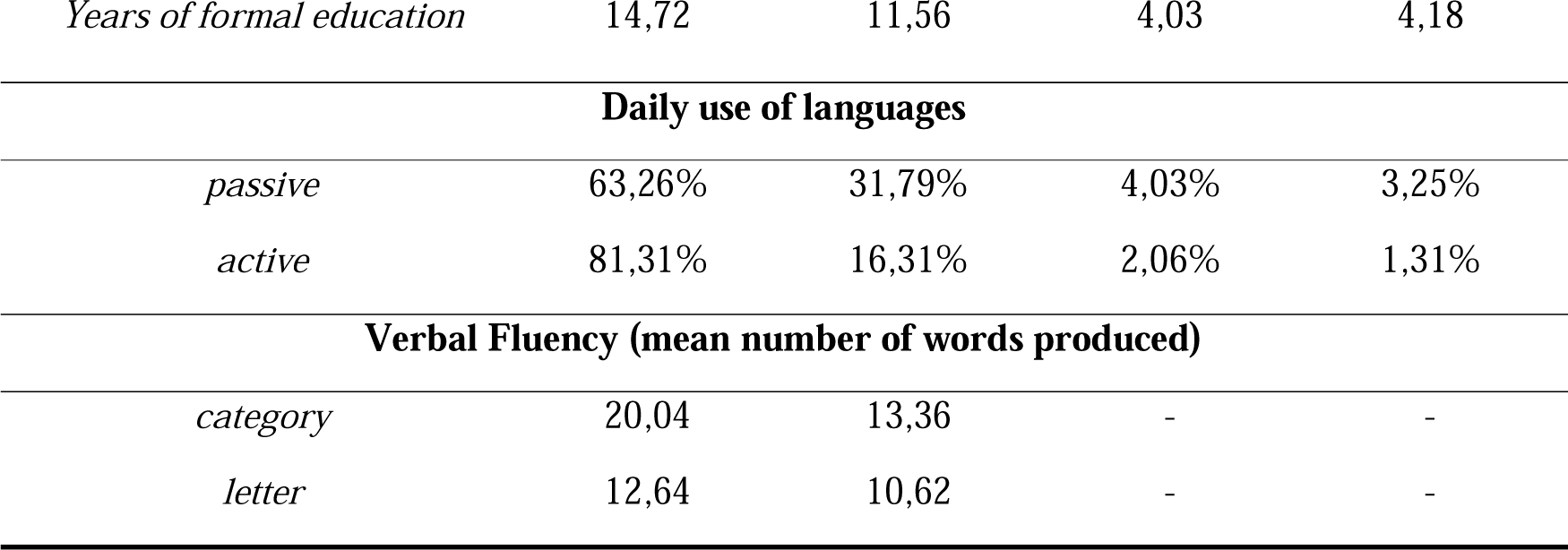
Language experience of participants. Information on self-rated proficiency, language learning and daily use of languages is given for all languages that participants declared to know. L1 always refers to Polish and L2 to English. L3 and L4 were a variety of different languages (incl. German, French, Italian, Spanish, Russian, Czech, Japanese, Korean, Norwegian, Latin and Esperanto). Not all the participants reported to know an L3 or L4.

Informed consent was obtained from all the study participants. The study was approved by the Ethics Committee of Jagiellonian University Institute of Psychology concerning experimental studies with human subjects and all methods were performed in accordance with the relevant guidelines and regulations of the Declaration of Helsinki.

### Study design

Each participant completed a series of behavioural tasks (for details see [79]) to assess their language proficiency as well as a detailed language questionnaire. Their results are summarized in Table 5. In the scanner, each subject completed a main task (picture naming in L2 and L1) and the Language and Multiple Demand network localizers. To keep the language context under control, the main task was split between two sessions: in one of them, subjects completed the picture naming task in L2 and in another, in L1. Several additional tasks were included in the experiment, however, they’re beyond the scope of this paper.

#### Main task: picture naming

In the main task, participants were asked to overtly name pictures of objects. To in order to minimize effects of language switching, which were not of theoretical interest in this study, participants named pictures in L1 and L2 on two separate sessions (on different days). The order of sessions was counterbalanced between participants. In each session, participants named one block of pictures corresponded to one functional run. Each block was composed of 55 coloured pictures representing objects (from the CLT database [81]). Pictures were not repeated between blocks. Each block started with a fixation (“+”) displayed on a screen for 7000 ms. Subsequently, each picture was displayed on a white background for 2000 ms and was followed by a fixation jittered at ITI ranging from 607 ms to 10144 ms (M = 3091 ms, SD = 2225 ms). The order of picture presentation and interstimulus intervals was optimized using optseq2 [82]. Ech block concluded with a fixation presented for 8000 ms. Overall, each block of the Picture Naming task lasted 295s.

#### Language localizer

Participants listened to fragments of Alice in Wonderland in L1 and distorted speech recordings in which it was impossible to understand what the speaker was saying [33]. The localizer contrast was based on the intact > distorted speech condition contrast. In each functional run, participants listened to 6 short passages of Alice in Wonderland in their native language (18s each) and 6 passages of distorted speech (18s each). Additionally, 4 fixation blocks (12s each) were included in each run. The total duration of each run was 264s. Each participant completed 2 runs of the task.

#### Multiple Demand localizer

Participants were asked to perform a spatial working memory task [23,31]. The localizer contrast was based on the hard > easy condition contrast. In each trial, participants saw four 3 x 4 grids and their task was to memorize the locations of black fields in each grid. In the easy condition, they had to memorize one location per grid had to be memorized and in the hard condition, they had to keep track of two locations per grid. At the end of each trial, participants were asked to choose a grid with all the memorized locations, from among two alternatives presented on the screen. Each trial started with a fixation displayed on a screen for 500 ms and was followed by four grids, each displayed for 1000 ms. After that, the choice task appeared on the screen until a response was given (for a max of 3750 ms) followed by feedback (right or wrong) displayed for 250 ms and a fixation which was displayed for 3750 ms minus the reaction time in the choice task. In each of the conditions – easy and hard – participants completed 4 trials (34s in total). Each run contained five easy condition blocks and five hard condition blocks. Additionally, six fixation blocks (16s) were included in each run. The order of experimental and fixation blocks was counterbalanced between participants and runs (four counterbalance lists were created and each participant completed two of them). The total duration of one run was 438s.

### MRI Data Acquisition

MRI data were acquired using a 3T scanner (Magnetom Skyra, Siemens) with a 64-channel head coil. High-resolution, whole-brain anatomical images were acquired using a T1-MPRAGE sequence (208 sagittal slices; voxel size 0.9×0.9×0.9 mm3; TR = 1800 ms, TE = 2.32 ms, flip angle = 8°). A gradient field map was acquired with a dual-echo gradient-echo sequence, matched spatially with fMRI scans (TE1 = 4.92 ms, TE2 = 7.38 ms, TR = 503 ms, flip angle = 60°). Functional T2*-weighted images were acquired using a whole-brain echo-planar (EPI) pulse sequence (50 axial slices, 3×3×3 mm isotropic voxels; TR = 1400 ms; TE = 27 ms; flip angle = 70°; MB acceleration factor 2; in-plane GRAPPA acceleration factor 2 and phase encoding A>>P) using 2D multiband echo-planar imaging (EPI) sequences from the Center for Magnetic Resonance Research (CMRR), University of Minnesota [83].

### Data preprocessing

All data were visually inspected for artifacts. The non-brain tissue was removed using the FSL Brain Extraction Tool [84]. We preprocessed the functional data, using FSL FEAT (version 6.0.0 [85]). The preprocessing included high-pass temporal filtering (100s), spatial smoothing (FWHM = 5 mm), distortion correction using fieldmap with FUGUE, co-registration, and normalization using FLIRT [86,87] and motion correction using MCFLIRT [86] with 6 rigid-body transformations. Functional images were first co-registered to their anatomical images and then registered to an MNI standard space (FSL’s 2mm standard brain template).

### Data analysis

Neuroimaging data used in these analyses is stored in an OpenNeuro repository (https://openneuro.org/datasets/ds004456). Data and code necessary to reproduce the analyses and plots reported in this paper is available in the OSF repository (https://osf.io/s85hk/?view_only=f37a4a291bfd49f690f97548db66e80c).

### Behavioral data analysis

Behavioral naming data recorded in the scanner during the picture naming task were transcribed and analyzed for naming latencies and accuracy. All answers corresponding to the dominant response and its synonyms were considered correct. Trials with no responses, inaudible responses and names unrelated to a given picture were classified as errors. We fitted a linear mixed-effect model to the naming latency data and a generalized mixed-effect linear model to the accuracy data (using the lmer() and glmer() functions from the lme4 for package [88] respectively). For both models we used the same formula:

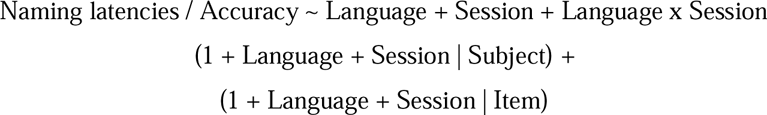

*Language* corresponded to a predictor with two levels: *L1* and *L2* and *Session* corresponded to a predictor with two levels: *first* and *second session*. Prior to the analysis the categorical predictors were deviation coded (L1 = −0.5; L2 = 0.5 and first session = −0.5; second session = 0.5). The analysis was performed using the lmer() function from the lme4 package. For each analysis, we first fitted a maximal model and then identified the best random effects’ structure [89]. Post-hoc pairwise comparisons were run using the lsmeans() function from the lsmeans package.

### Neuroimaging data analyses

In the 1^st^ level statistical analysis, a double-gamma hemodynamic response function was used to model the BOLD signal corresponding to each event (trial in the main task and blocks in the localizer tasks). The estimates of motion obtained with MCFLIRT were included in the 1^st^ level GLM as nuisance covariates. We performed two types of analyses of the picture naming data: the localizer-based analyses and the whole-brain analysis.

#### Localizer-based analyses

Two functional runs of each localizer task were combined in a second-level analysis for each participant using an FSL fixed-effect model. To ensure better comparability of our findings with previous studies, as group-level localizer masks we used sets of functional parcels established and validated by previous studies instead of creating the group-level parcels based on our data (twelve language parcels [32]; twenty MD parcels [23]). The group-level parcels used in this analysis are available in the OSF repository (https://osf.io/s85hk/?view_only=f37a4a291bfd49f690f97548db66e80c). These group-level partitions were intersected with subject-specific data yielding a set of subject-specific group-constraint ROIs. Finally, we selected the top 10% most responsive voxels within each parcel based on the *z*-value map for a given localizer contrast (Language: speech > degraded speech, MD: hard > easy visual working memory task) and we binarized the obtained ROIs. At the single subject level, the language and MD fROIs showed a minimal overlap (mean = 0,172%, SD = 0,26%). While the mean overlap of the individual-level fROIs used in this study has been constrained by the group-level parcels, the overlap of the whole-brain maps for the language and MD localizer contrast, computed for each participant separately, thresholded at 10% of the most active voxels, was also very small (mean = 6,58%, SD = 4,67%) showing that the two localizer tasks identify independent functional systems in the brain (in line with). Data used to calculate the overlap statistics is available in the OSF repository. We then used the functionally-defined ROIs for the language and MD networks to extract parameter estimates corresponding to % signal change in the two critical conditions of the main task: naming pictures in L1 and L2. Parameter estimates were extracted from the 1^st^ level analyses for each participant separately. This was done using an FSL’s FEATquery tool (http://www.FMRIb.ox.ac.uk/fsl/feat5/featquery.html). Further analyses were performed on the extracted parameter estimates in R (version: 4.0.2). Following an approach that proposes to model activity within a functional network on the network level, instead of relying on separate analysis for each ROI we fitted one linear mixed-effect model for each functional network according to the following formula:

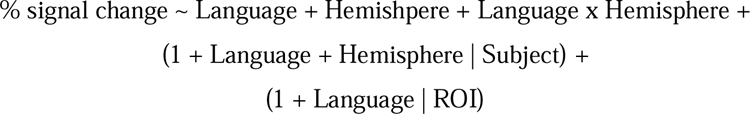

In each model, the fixed effect of Language corresponded to a predictor with two levels: *L1* and *L2* and the effect of *Hemisphere* corresponded to a predictor with two levels: *left hemisphere* and *right hemisphere*. Prior to the analysis this categorical predictor was deviation coded (L1 = −0.5; L2 = 0.5 and left hemisphere = −0.5; right hemisphere = 0.5).

To further explore our results, responses to the language and MD localizer tasks conditions were analyzed using the following linear mixed-effect model fitted separately for each ROI:

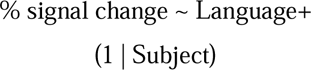

Prior to the analysis, the categorical predictor of condition (intact or degraded speech for the language localizer task and hard or easy visual WM for the MD localizer task) was deviation coded. The resulting *p*-values were corrected for multiple comparisons using the FDR correction.

To provide a more direct link between brain activation and the difficulty of speech production, we have also analysed the relationship between brain response in the Language and MD networks with mean reaction times (RT) in a picture naming task. To this aim, we fitted one linear mixed-effect model for each functional network according to the following formula:

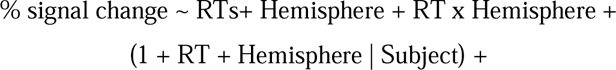

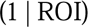

In this analysis, we used a mean RT score for each subject. The predictor of mean RTs (in seconds) was demanded prior to the analysis.

Finally, to assess the degree to which differences in brain response to speech production in L1 and L2 are modulated by individual differences between participants, concerning mostly their language experience, we tested how the response to L1 vs L2 in the left and right hemisphere is modulated by variables describing individual differences between participants. The following variables were used as predictors: years of education (counted from the 1^st^ year of the primary school), fluid intelligence (measured with a shortened version of the Raven test), proficiency in L2 (measured with LexTale), age of acquisition of L2, and the percent of active and passive daily use of L1 (in respect to other languages known by each participant). To this aim, we fitted one linear mixed-effect model for each functional network according to the following formula:

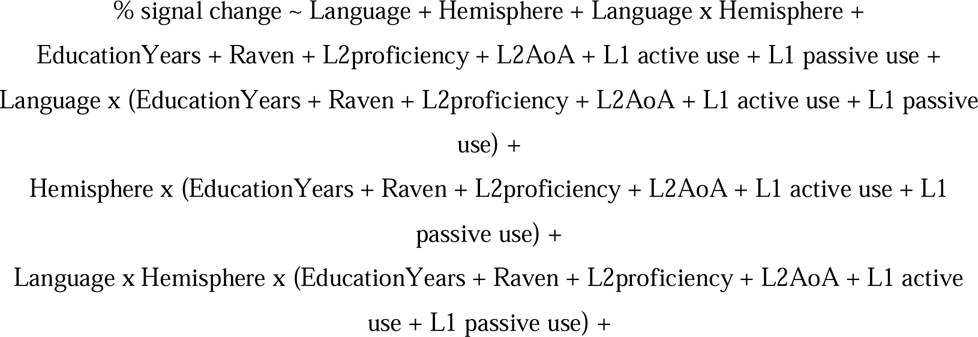

Prior to these analyses, the categorical predictors were deviation-coded and the continuous predictors we demeaned.

All the analyses was performed using the lmer() function from the lme4 package [88]. For each functional network, we first fitted a maximal model and then identified the best random effects’ structure [89].

#### Bilingual language control network analysis

To evaluate language and domain-general selective responses within the Bilingual Language Control network, we used a similar approach as in the localizer-based analyses. First, we selected a set of ROIs corresponding to the Bilingual Language Control network, as described by Abutalebi and Green [15]. These included the left inferior frontal gyrus (IFG) – pars opercularis and pars triangularis, left middle frontal gyrus (MFG), left anterior cingulate cortex, left pre-supplementary motor area (pre-SMA), left angular gyrus, left thalamus, putamen and caudate as well as right IFG – pars opercularis and triangularis and right angular gyrus. The ROIs were derived from the FSL Harvard-Oxford Cortical and Subcortical atlases. As the atlases provide probabilistic masks, the selected ROIs were thresholded at 30% and binarized prior to further analyses. The pre-SMA is not available as a separate mask within the Harvard-Oxford Cortical atlas, hence it was created by splitting the Supplementary Motor Cortex ROI into anterior and posterior parts along y=0. Final set of masks used for this analysis is available at OSF (https://osf.io/s85hk/?view_only=f37a4a291bfd49f690f97548db66e80c). To create subject-specific language and domain-general selective masks within the BLC ROIs, each of the anatomically-defined ROI was intersected with activation map for each subject for corresponding to the (i) language localizer contrast, i.e., intact > degraded speech; and (ii) the MD localizer contrast, i.e., hard vWM > easy vWM. Subsequently, we selected top 10% of the most responsive voxels for each contrast and we binarized the obtained maps to create subject-specific functional ROIs (fROIs) within each of the anatomically-defined BLC ROI.

Following that, response to L1 and L2 speech production in a picture naming task were extracted from the language- and domain-general selective fROIs using the same approach as described in the localizer-based analyses.

To assess the voxel overlap between the two networks, individual subject parcels for language and domain-general fROIs within each BLC network ROI were overlaid in a common space and a number of voxels overlapping between the two masks was calculated. Following that, the proportion of the overlapping voxels in each BLC network ROI was calculated using the following formula:

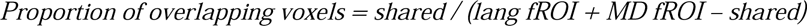

Where shared corresponds to the number of voxels shared between the two fROIs and lang fROI and MD fROI corresponds to the total number of voxels in fROI defined using the language and MD localizers.

## Acknowledgements

We would like to thank all members of the Psychology of Language and Bilingualism Lab who contributed to this project by collecting and coding the data and to all participants who took part in the study. We are very grateful to Evelina Fedorenko for inspiration and insightful comments on the design and results.

## Author contributions

AW, JS, MD, MS, and ZW contributed to the conception, design and methodology of the study. AW and AD collected the data and AW designed and conducted data analysis. AW, JS, MD and ZW interpreted the analyses. ZW was responsible for project supervision. AW, JS and ZW wrote the first draft of the manuscript and AD and MD provided critical review and revision the text. All authors approved the final version for publication.

## Data availability statement

The raw neuroimaging that served as a basis for analyses in this paper along with the data on participants’ language experience is available at https://openneuro.org/datasets/ds004456.

Data and code necessary to reproduce the presented ROI and behavioural analyses is available at the https://osf.io/s85hk/?view_only=f37a4a291bfd49f690f97548db66e80c.

## Additional Information

Authors declare no competing interests.

