## Supplementary Materials for "Domain-general and language-specific contributions to speech production in a second language: an fMRI study using functional localizers"

### 30-060 Kraków

### 1. Whole brain analysis

**Statistical analyses methods:**

For the whole-brain analysis, functional runs corresponding to naming in L1 and L2 were combined in a second-level analysis using an FSL fixed-effect model which allowed to create a single-subject contrast between the conditions of interest. Subsequently, a group-level paired t-test was run using an FSL mixed effects model analysis (FLAME 1+2). Group activations were thresholded at z>3.1 and corrected for multiple comparisons using FSL’s cluster-based interference to *p*<0.05. We tested two contrasts: L2 > L1 and L1 > L2.

**Results:**

The whole-brain analysis revealed that naming in L2 compared to naming in L1 (L2>L1) elicited increased neural activity in a number of clusters located in the left occipital cortex extending to the left cuneus, left thalamus and the right and left caudate nuclei, left lingual gyrus, right supplementary motor area, left superior frontal gyrus and left frontal pole extending, left inferior frontal gyrus (pars opercularis) extending to the left precentral gyrus, right precentral gyrus and right superior frontal gyrus. The reverse contrast (L1>L2) revealed that naming in L1 compared to naming in L2 elicited increased neural activity in the left angular/supramarginal gyrus, right posterior superior temporal gyrus, left putamen, right operculum, extending to the Heschl’s gyrus, left anterior middle temporal gyrus and the right putamen. The results of the comparison between languages are summarized in **Table S1 and Figure S1.**

**Table S1. Whole-brain comparison between speech production in L2 and L1.** Significant clusters of activations corrected for multiple comparison at the voxel level (*z* > 3.1) and cluster level (*p* < .05).

|  | **Volume** | | | | **Peak** | | |
| --- | --- | --- | --- | --- | --- | --- | --- |
|  | **Hemisphere** | **n voxels** | ***p* – value** | **­­*z* – score** | **x** | **y** | **z** |
| **L2 > L1** | | | | | | | |
| Precuneous / OccipitalPole | Left | 1798 | 0,000 | 5,00 | 4 | -94 | 16 |
| Thalamus | Left | 1402 | 0,000 | 5,43 | -2 | -4 | 10 |
| Caudate | Right |  |  | 4,45 | 10 | 4 | 8 |
| Caudate | Left |  |  | 4,43 | -10 | 6 | 10 |
| Lingual Gyrus | Left | 831 | 0,000 | 4,83 | -8 | -72 | -6 |
| Paracingulate Gyrus (SMA) | Right | 768 | 0,000 | 5,04 | 4 | 2 | 72 |
| Superior Frontal gyrus | Left | 543 | 0,000 | 4,45 | -22 | -8 | 72 |
| Postcentral Gyrus (Superior Parietal Lobule) | Left | 445 | 0,000 | 4,68 | -40 | -34 | 56 |
| Frontal Pole | Left | 352 | 0,000 | 5,00 | -44 | 44 | 14 |
| Inferior Frontal Gyrus (opercularis) / Precentral Gyrus | Left | 297 | 0,000 | 4,48 | -50 | 14 | 32 |
| Precentral Gyrus | Right | 118 | 0,023 | 4,11 | 4 | -26 | 76 |
| Superior Frontal Gyrus | Right | 103 | 0,042 | 4,23 | 24 | 2 | 72 |
| **L1 > L2** | | | | | | | |
| Angular / Supramarginal Gyrus | Left | 717 | 0,000 | 5,04 | -58 | -58 | 26 |
| Superior Temporal Gyrus (posterior part) | Right | 286 | 0,000 | 4,67 | 58 | -24 | 2 |
| Putamen | Left | 136 | 0,012 | 4,52 | -28 | -6 | 6 |
| Operculum | Right | 135 | 0,012 | 3,97 | 58 | 0 | 0 |
| Middle Temporal Gyrus (anterior part) | Left | 113 | 0,028 | 4,31 | -50 | -4 | -30 |
| Putamen | Right | 100 | 0,048 | 4,22 | 30 | -14 | 4 |

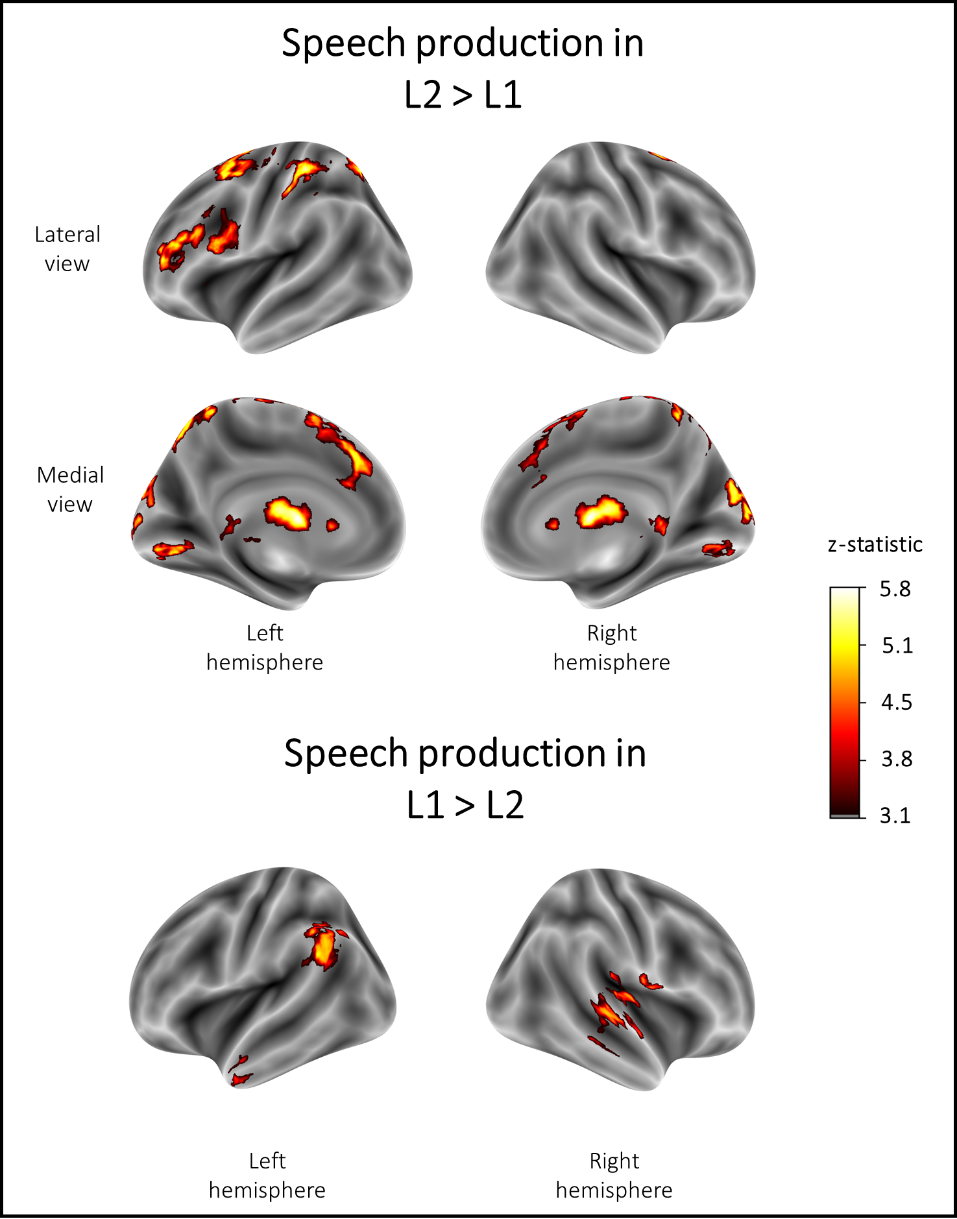

**Figure S1. Whole brain comparison between speech production in L2 > L1.** All presented activations are corrected at the voxel level (*z* > 3.1) and cluster level (*p* < 0.05).

| Influence of naming latencies in brain activation in the language and MD networks  To provide a more direct link between brain activation and the difficulty of speech production, we have also analyzed the relationship between brain response in the Language and MD networks with mean reaction times (RT) in a picture naming task. To this aim, we fitted one linear mixed-effect model for each functional network according to the following formula:  % signal change ~ RTs+ Hemisphere + RT x Hemisphere +  (1 + RT + Hemisphere \| Subject) +  (1 \| ROI)  In this analysis, we used a mean RT score for each subject. The predictor of mean RTs (in seconds) was demanded prior to the analysis. Table S2 presents the results of these analyses.  Responses to the language and MD localizer tasks conditions were subsequently analyzed using the following linear mixed-effect model fitted separately for each ROI:  % signal change ~ RT+  (1 \| Subject)  The resulting p-values were corrected for multiple comparisons using the FDR correction. Tables S3 and S4 present the results of these analyses for the MD and language fROIs, respectively.  **Table S2. Estimated brain response in language and MD networks predicted by naming latencies.** The table presents results of linear mixed models predicting the % BOLD signal change in the two networks predicted by mean reaction times for L1 and L2 picture naming for each subject. | | | | | |
| --- | --- | --- | --- | --- | --- |
|  | **Language network** | | | | |
| *Predictors* | *Estimates* | *std. Error* | *CI* | *t-*value | *p* – value |
| Intercept | 0.57 | 0.11 | 0.34  0.79 | 5.00 | **<0.001** |
| Reaction Times | -0.06 | 0.20 | -0.45 0.32 | -0.33 | 0.744 |
| Hemisphere (left vs right) | -0.13 | 0.21 | -0.55  0.28 | -0.63 | 0.529 |
| Language x Hemisphere | -0.29 | 0.20 | -0.67  0.10 | -1.47 | 0.142 |
|  | **MD network** | | | | |
| *Predictors* | *Estimates* | *std. Error* | *CI* | *Statistic* | *p - value* |
| Intercept | 0.48 | 0.07 | 0.35  0.61 | 7.32 | **<0.001** |
| Reaction Times | 0.76 | 0.22 | 0.33  1.19 | 3.49 | **<0.001** |
| Hemisphere (left vs right) | -0.20 | 0.09 | -0.38  -0.02 | -2.19 | **0.029** |
| Language x Hemisphere | -0.61 | 0.14 | -0.90  -0.33 | -4.26 | **<0.001** |

**Table S3** **Results of linear mixed-effect models fitted for each functional ROI within the language network.** Presented estimates correspond to the of mean naming latencies for each language.

| **Language network** | | | | | | | | | |
| --- | --- | --- | --- | --- | --- | --- | --- | --- | --- |
| **Region of interest** | | | **Fixed effect of Reaction Times**  (mean for L2 and L1) | | | | | | |
| **Hemisphere** | **ROI Label** | **Estimate** | | **Standard  Error** | ***t* - value** | **Effect size *d*** | ***p*-value** | | |
|  |  |  |  |  |  |  | ***uncorrected*** | | ***FDR- corrected*** |
| left | IFGorb | 0,22 | | 0,32 | 0,69 | 0,18 | 0,494 | 0,593 | |
| left | IFGoperc | 0,78 | | 0,25 | 3,17 | 0,89 | 0,003 | 0,031* | |
| left | MFG | 0,55 | | 0,39 | 1,39 | 0,41 | 0,170 | 0,308 | |
| left | antTemp | -0,27 | | 0,25 | -1,09 | -0,29 | 0,279 | 0,419 | |
| left | postTemp | -0,35 | | 0,20 | -1,73 | -0,49 | 0,090 | 0,308 | |
| left | AngG | 0,44 | | 0,29 | 1,53 | 0,40 | 0,131 | 0,308 | |
| right | IFGorb | -0,38 | | 0,28 | -1,36 | -0,36 | 0,180 | 0,308 | |
| right | IFGoperc | 0,04 | | 0,18 | 0,22 | 0,06 | 0,828 | 0,904 | |
| right | MFG | -0,28 | | 0,38 | -0,73 | -0,21 | 0,467 | 0,593 | |
| right | antTemp | -0,30 | | 0,21 | -1,44 | -0,38 | 0,155 | 0,308 | |
| right | postTemp | -0,49 | | 0,24 | -2,10 | -0,53 | 0,040 | 0,238 | |
| right | AngG | -0,01 | | 0,27 | -0,05 | -0,01 | 0,961 | 0,961 | |

**Table S4.** **Results of linear mixed-effect models fitted for each functional ROI within the MD network.** Presented estimates correspond to the effect of mean naming latencies for each language.

| **Multiple Demand network** | | | | | | | |
| --- | --- | --- | --- | --- | --- | --- | --- |
| **Region of interest** | | **Fixed effect of Reaction Times**  (mean for L2 and L1) | | | | | |
| **Hemisphere** | **ROI Label** | **Estimate** | **Standard  Error** | ***t* - value** | **Effect size *d*** | ***p*-value** | |
|  |  |  |  |  |  | ***uncorrected*** | ***FDR-corrected*** |
| left | midFrontalOrb | 1,00 | 0,33 | 3,00 | 0,80 | 0,004 | 0,011* |
| left | midFrontal | 1,07 | 0,30 | 3,60 | 0,96 | 0,001 | 0,003* |
| left | insula | 0,55 | 0,23 | 2,38 | 0,62 | 0,021 | 0,037* |
| left | IFGoper | 1,09 | 0,26 | 4,26 | 1,22 | 0,000 | 0,001* |
| left | PrecG | 0,79 | 0,28 | 2,85 | 0,79 | 0,006 | 0,014* |
| left | supFrontal | 1,08 | 0,27 | 3,97 | 1,15 | 0,000 | 0,001* |
| left | SMA | 0,91 | 0,23 | 3,95 | 1,09 | 0,000 | 0,001* |
| left | antParietal | 0,66 | 0,24 | 2,82 | 0,74 | 0,007 | 0,014* |
| left | midParietal | 0,36 | 0,37 | 0,96 | 0,28 | 0,341 | 0,487 |
| left | postParietal | 1,50 | 0,35 | 4,24 | 1,21 | 0,000 | 0,001* |
| right | midFrontalOrb | 0,39 | 0,23 | 1,65 | 0,44 | 0,105 | 0,161 |
| right | midFrontal | 0,16 | 0,20 | 0,81 | 0,21 | 0,421 | 0,526 |
| right | insula | 0,56 | 0,18 | 3,11 | 0,80 | 0,003 | 0,010* |
| right | IFGoper | -0,09 | 0,21 | -0,42 | -0,12 | 0,673 | 0,748 |
| right | PrecG | -0,07 | 0,22 | -0,34 | -0,09 | 0,737 | 0,766 |
| right | supFrontal | 0,17 | 0,25 | 0,67 | 0,17 | 0,509 | 0,598 |
| right | SMA | 0,61 | 0,22 | 2,81 | 0,77 | 0,007 | 0,014* |
| right | antParietal | -0,19 | 0,23 | -0,83 | -0,22 | 0,408 | 0,526 |
| right | midParietal | 0,09 | 0,32 | 0,30 | 0,08 | 0,766 | 0,766 |
| right | postParietal | 0,68 | 0,37 | 1,85 | 0,51 | 0,070 | 0,117 |

### Relationship between individual differences and language experience on brain activation in response to L1 and L2 in the langauge and MD networks

To assess the degree to which differences in brain response to speech production in L1 and L2 are modulated by individual differences between participants, concerning mostly their language experience, we tested how the response to L1 vs L2 in the left and right hemisphere is modulated by variables describing individual differences between participants. The following variables were used as predictors: years of education (counted from the 1^st^ year of the primary school), fluid intelligence (measured with a shortened version of the Raven test), proficiency in L2 (measured with LexTale), age of acquisition of L2, and the percent of active and passive daily use of L1 (in respect to other languages known by each participant). To this aim, we fitted one linear mixed-effect model for each functional network according to the following formula:

% signal change ~

Language + Hemishpere + Language x Hemisphere +

EducationYears + Raven + L2proficiency + L2AoA + L1 active use + L1 passive use +

Language x (EducationYears + Raven + L2proficiency + L2AoA + L1 active use + L1 passive use) +

Hemisphere x (EducationYears + Raven + L2proficiency + L2AoA + L1 active use + L1 passive use) +

Language x Hemisphere x (EducationYears + Raven + L2proficiency + L2AoA + L1 active use + L1 passive use) +

Prior to these analyses, the categorical predictors were deviation-coded and the continuous predictors we demeaned. Detailed results of these analyses are presented in Tables S5 and S6. The relationship between the proficiency, language, and hemisphere in the MD network (the only significant effect we found in the analysis of the influence of individual differences and language experience with the neuroimaging results) is presented in Figure S2.)

**Table S5.** **Estimated brain response in language network predicted by language, hemisphere and individual differences and language experience of participants.**

|  | **Language network**  Effects of language, hemisphere, and individual differences | | | | |
| --- | --- | --- | --- | --- | --- |
| *Predictors* | *Estimates* | *std. Error* | *CI* | *t-*value | *p* – value |
| (Intercept) | 0.56 | 0.11 | 0.34  0.78 | 4.95 | **<0.001** |
| Language (L2 vs L1) | -0.02 | 0.04 | -0.09  0.05 | -0.64 | 0.521 |
| Hemisphere (left vs right) | -0.13 | 0.21 | -0.54  0.28 | -0.61 | 0.541 |
| Language × Hemisphere | -0.12 | 0.04 | -0.21  -0.04 | -2.81 | **0.005*** |
| Education Years | 0.05 | 0.02 | 0.01  0.08 | 2.69 | **0.007*** |
| Raven | 0.20 | 0.31 | -0.40  0.81 | 0.65 | 0.513 |
| L2 proficiency | -0.54 | 0.53 | -1.57  0.49 | -1.02 | 0.306 |
| L2 AoA | 0.00 | 0.02 | -0.04  0.03 | -0.20 | 0.845 |
| %L1 active use | 0.00 | 0.00 | 0.00  0.01 | 0.85 | 0.397 |
| %L1 passive use | 0.00 | 0.00 | -0.01  0.00 | -0.67 | 0.504 |
| Edu. Years × Language | 0.00 | 0.02 | -0.03  0.03 | -0.22 | 0.828 |
| Raven × Language | -0.02 | 0.27 | -0.55  0.51 | -0.08 | 0.936 |
| L2 prof × Language | 0.02 | 0.46 | -0.88  0.92 | 0.05 | 0.963 |
| L2 AoA × Language | 0.02 | 0.02 | -0.02  0.05 | 0.92 | 0.356 |
| %L1 active use × Language | 0.00 | 0.00 | -0.01  0.00 | -1.19 | 0.234 |
| %L1 passive use × Language | 0.00 | 0.00 | 0.00  0.01 | 0.12 | 0.902 |
| Edu. Years × Hemisphere | -0.01 | 0.01 | -0.03  0.02 | -0.54 | 0.592 |
| Raven × Hemisphere | 0.18 | 0.20 | -0.21  0.56 | 0.90 | 0.368 |
| L2 prof × Hemisphere | 0.32 | 0.33 | -0.33  0.97 | 0.96 | 0.335 |
| L2 AoA × Hemisphere | 0.00 | 0.01 | -0.03  0.02 | -0.33 | 0.742 |
| %L1 active use × Hemisphere | 0.00 | 0.00 | -0.01  0.00 | -0.65 | 0.516 |
| %L1 passive use × Hemisphere | 0.00 | 0.00 | 0.00  0.00 | 0.47 | 0.635 |
| Education Years ×  Language × Hemisphere | 0.02 | 0.02 | -0.01  0.06 | 1.19 | 0.234 |
| Raven ×  Language × Hemisphere | -0.07 | 0.33 | -0.72  0.58 | -0.20 | 0.841 |
| L2 proficiency × Language × Hemisphere | 0.90 | 0.56 | -0.20  2.01 | 1.60 | 0.110 |
| L2 AoA ×  Language × Hemisphere | 0.00 | 0.02 | -0.04  0.04 | -0.20 | 0.838 |
| %L1 active use ×  Language × Hemisphere | 0.00 | 0.00 | -0.01  0.01 | 0.06 | 0.954 |
| %L1 passive use ×  Language × Hemisphere | 0.00 | 0.00 | 0.00  0.01 | 1.21 | 0.228 |

**Table S5.** **Estimated brain response in MD network predicted by language, hemisphere and individual differences and language experience of participants.**

|  | **MD network**  Effects of language, hemisphere, and individual differences | | | | |
| --- | --- | --- | --- | --- | --- |
| *Predictors* | *Estimates* | *std. Error* | *CI* | *t-*value | *p* – value |
| (Intercept) | 0.46 | 0.06 | 0.35  0.57 | 7.88 | **<0.001** |
| Language | 0.11 | 0.03 | 0.04  0.18 | 3.22 | **0.001*** |
| Hemisphere | -0.19 | 0.09 | -0.37  -0.02 | -2.14 | **0.033*** |
| Language × Hemisphere | -0.15 | 0.03 | -0.21  -0.09 | -5.14 | **<0.001*** |
| Education Years | 0.06 | 0.02 | 0.02  0.09 | 3.45 | **0.001*** |
| Raven | 0.48 | 0.29 | -0.09  1.04 | 1.66 | 0.097 |
| L2 proficiency | -0.55 | 0.49 | -1.51  0.41 | -1.12 | 0.264 |
| L2 AoA | 0.03 | 0.02 | -0.01  0.06 | 1.62 | 0.104 |
| %L1 active use | 0.00 | 0.00 | 0.00  0.01 | 1.60 | 0.109 |
| %L1 passive use | 0.00 | 0.00 | -0.01  0.00 | -0.87 | 0.385 |
| Edu. Years × Language | 0.01 | 0.01 | -0.02  0.04 | 0.63 | 0.529 |
| Raven × Language | -0.19 | 0.25 | -0.69  0.30 | -0.77 | 0.444 |
| L2 prof × Language | -0.61 | 0.43 | -1.46  0.24 | -1.41 | 0.160 |
| L2 AoA × Language | -0.01 | 0.02 | -0.04  0.02 | -0.51 | 0.610 |
| %L1 active use × Language | 0.00 | 0.00 | -0.01  0.00 | -0.43 | 0.669 |
| %L1 passive use × Language | 0.00 | 0.00 | -0.01  0.00 | -0.64 | 0.525 |
| Edu. Years × Hemisphere | -0.01 | 0.01 | -0.03  0.01 | -1.05 | 0.296 |
| Raven × Hemisphere | -0.28 | 0.18 | -0.63  0.07 | -1.57 | 0.116 |
| L2 prof × Hemisphere | 0.37 | 0.30 | -0.22  0.96 | 1.22 | 0.223 |
| L2 AoA × Hemisphere | -0.01 | 0.01 | -0.04  0.01 | -1.32 | 0.186 |
| %L1 active use × Hemisphere | 0.00 | 0.00 | 0.00  0.00 | 0.07 | 0.944 |
| %L1 passive use × Hemisphere | 0.00 | 0.00 | 0.00  0.00 | -0.15 | 0.880 |
| Education Years ×  Language × Hemisphere | 0.01 | 0.01 | -0.02  0.03 | 0.47 | 0.638 |
| Raven ×  Language × Hemisphere | -0.41 | 0.22 | -0.84  0.02 | -1.86 | 0.063 |
| L2 proficiency × Language × Hemisphere | 0.79 | 0.38 | 0.05  1.52 | 2.09 | **0.037*** |
| L2 AoA ×  Language × Hemisphere | -0.02 | 0.01 | -0.04  0.01 | -1.21 | 0.226 |
| %L1 active use ×  Language × Hemisphere | 0.00 | 0.00 | 0.00  0.01 | 0.88 | 0.381 |
| %L1 passive use ×  Language × Hemisphere | 0.00 | 0.00 | 0.00  0.01 | 0.94 | 0.348 |

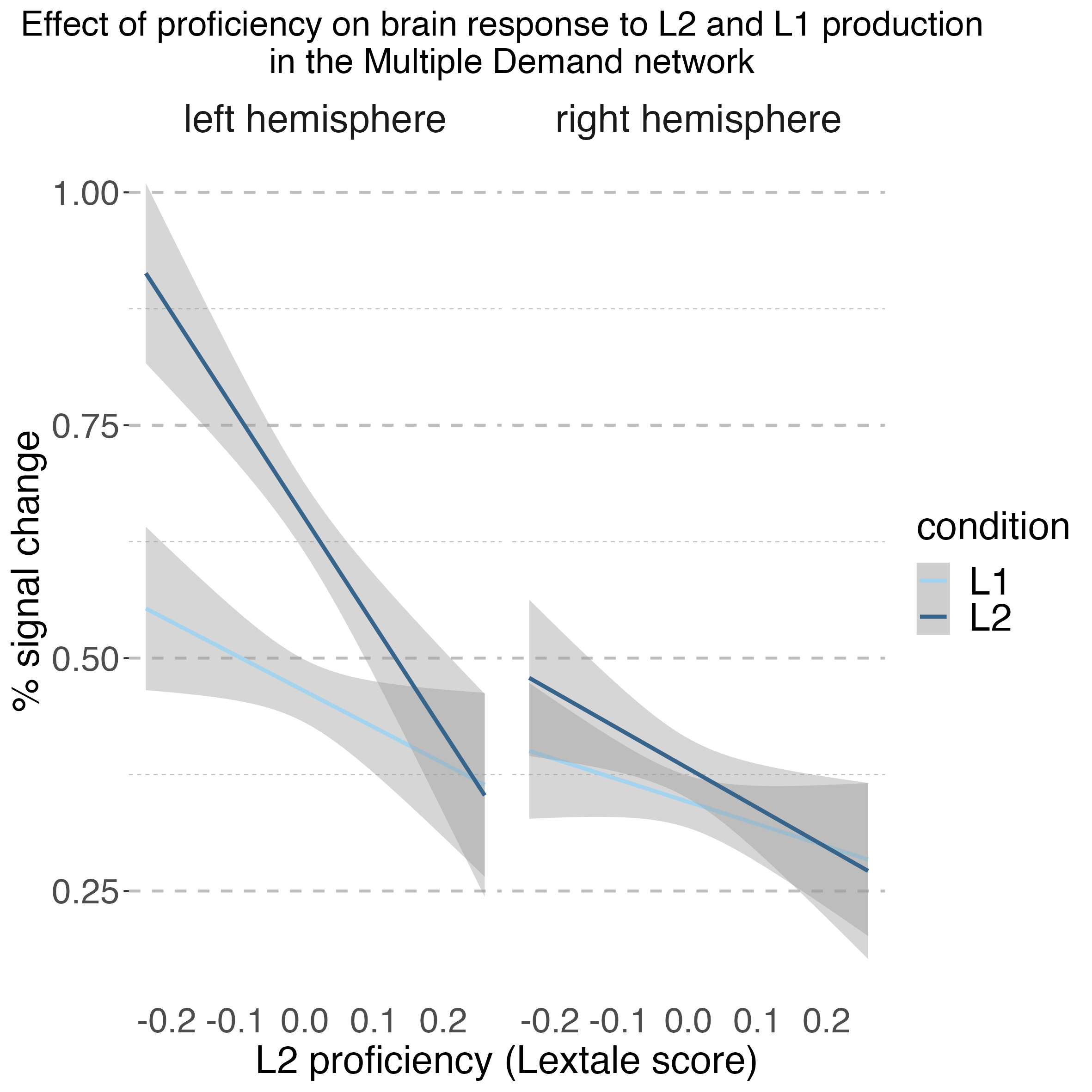

**Figure S2. Relationship between language (L2 vs L1) and proficiency level in the right and left hemisphere within the MD network**. This figure represents the effect of an interaction between L2 proficiency, language, and hemisphere reported in Table S5.

### BLC ROIs: selectivity to the language and MD localizer tasks

To evaluate the selectivity of each fROI to its constituting localizer task, we extracted the parameter estimates corresponding to the two localizer tasks from the language and domain-general fROIs within each BLC ROI. To avoid circularity, we used an across-run cross-validation procedure to extract the estimates corresponding to the localizer task from the fROI defined using the same task: we defined the functional parcels based on the first functional run and estimated responses to the localizer contrast on the second run and repeated this procedure using the second run to define the functional parcels and the first run to extract response estimates (see Fedorenko et al., 2010). Responses to the language and MD localizer tasks conditions were subsequently analyzed using the following linear mixed-effect model fitted separately for each ROI:

% signal change ~ Language +

(1 | Subject)

Prior to the analysis, the categorical predictor of condition (intact or degraded speech for the language localizer task and hard or easy visual WM for the MD localizer task) was deviation coded. The resulting p-values were corrected for multiple comparisons using the FDR correction.

**Table S6.** **Results of linear mixed-effect models fitted for the localizer tasks in each BLC ROI.** Presented estimates correspond to the effect of the localizer tasks conditions (intact vs degraded in the langue localizer and hard vs easy visual WM for the MD localizer). ROI labels correspond to anatomical labels used in Figure 3 in the manuscript.

| **BLC ROI**  **(anatomical)** | **fROI**  **(localizer)** | **Contrast** | **Estimates** | **Standard  Error** | **t-value** | **Effect size *d*** | ***p*-value** | |
| --- | --- | --- | --- | --- | --- | --- | --- | --- |
|  |  |  |  |  |  |  | ***uncorrected*** | ***FDR- corrected*** |
| LeftACC | Language | intact > degraded | -0,04 | 0,02 | -1,76 | -0,32 | 0,081 | 0,108 |
|  |  | hard > easy vWM | 0,00 | 0,02 | -0,05 | -0,01 | 0,957 | 0,957 |
|  | MD | intact > degraded | -0,05 | 0,03 | -1,66 | -0,30 | 0,099 | 0,347 |
|  |  | hard > easy vWM | 0,12 | 0,04 | 3,15 | 0,57 | 0,002 | 0,002* |
| LeftAngularG | Language | intact > degraded | 0,42 | 0,06 | 7,25 | 1,31 | 0,000 | 0,000* |
|  |  | hard > easy vWM | -0,07 | 0,04 | -1,52 | -0,28 | 0,131 | 0,313 |
|  | MD | intact > degraded | -0,12 | 0,05 | -2,62 | -0,47 | 0,010 | 0,119 |
|  |  | hard > easy vWM | 0,27 | 0,06 | 4,18 | 0,76 | 0,000 | 0,000* |
| LeftCaudate | Language | intact > degraded | 0,02 | 0,02 | 1,06 | 0,19 | 0,290 | 0,316 |
|  |  | hard > easy vWM | 0,07 | 0,03 | 2,39 | 0,43 | 0,018 | 0,073 |
|  | MD | intact > degraded | 0,02 | 0,03 | 0,79 | 0,14 | 0,430 | 0,737 |
|  |  | hard > easy vWM | 0,12 | 0,03 | 3,73 | 0,67 | 0,000 | 0,000* |
| LeftIFGoper | Language | intact > degraded | 0,32 | 0,04 | 7,04 | 1,28 | 0,000 | 0,000* |
|  |  | hard > easy vWM | -0,01 | 0,04 | -0,20 | -0,04 | 0,845 | 0,921 |
|  | MD | intact > degraded | -0,02 | 0,04 | -0,39 | -0,07 | 0,696 | 0,835 |
|  |  | hard > easy vWM | 0,30 | 0,06 | 4,85 | 0,88 | 0,000 | 0,000* |
| LeftIFGtr | Language | intact > degraded | 0,33 | 0,05 | 6,81 | 1,23 | 0,000 | 0,000* |
|  |  | hard > easy vWM | -0,07 | 0,04 | -1,75 | -0,32 | 0,083 | 0,248 |
|  | MD | intact > degraded | -0,01 | 0,04 | -0,22 | -0,04 | 0,829 | 0,866 |
|  |  | hard > easy vWM | 0,20 | 0,05 | 3,70 | 0,67 | 0,000 | 0,000* |
| LeftMFG | Language | intact > degraded | 0,20 | 0,04 | 4,86 | 0,88 | 0,000 | 0,000* |
|  |  | hard > easy vWM | 0,14 | 0,05 | 2,77 | 0,50 | 0,006 | 0,073 |
|  | MD | intact > degraded | -0,06 | 0,04 | -1,51 | -0,27 | 0,133 | 0,347 |
|  |  | hard > easy vWM | 0,45 | 0,06 | 6,98 | 1,26 | 0,000 | 0,000* |
| LeftpreSMC | Language | intact > degraded | 0,13 | 0,04 | 3,72 | 0,67 | 0,000 | 0,001* |
|  |  | hard > easy vWM | 0,03 | 0,04 | 0,79 | 0,14 | 0,429 | 0,706 |
|  | MD | intact > degraded | 0,05 | 0,04 | 1,44 | 0,26 | 0,152 | 0,347 |
|  |  | hard > easy vWM | 0,16 | 0,06 | 2,66 | 0,48 | 0,009 | 0,010* |
| LeftPutamen | Language | intact > degraded | 0,04 | 0,02 | 2,15 | 0,39 | 0,034 | 0,050 |
|  |  | hard > easy vWM | 0,01 | 0,02 | 0,56 | 0,10 | 0,575 | 0,706 |
|  | MD | intact > degraded | 0,03 | 0,02 | 1,37 | 0,25 | 0,173 | 0,347 |
|  |  | hard > easy vWM | 0,04 | 0,02 | 1,89 | 0,34 | 0,061 | 0,061 |
| LeftThalamus | Language | intact > degraded | 0,02 | 0,03 | 0,85 | 0,15 | 0,399 | 0,399 |
|  |  | hard > easy vWM | 0,09 | 0,04 | 2,42 | 0,44 | 0,017 | 0,073 |
|  | MD | intact > degraded | 0,01 | 0,03 | 0,50 | 0,09 | 0,618 | 0,824 |
|  |  | hard > easy vWM | 0,13 | 0,04 | 3,55 | 0,64 | 0,001 | 0,001* |
| RightAngularG | Language | intact > degraded | 0,07 | 0,04 | 1,63 | 0,29 | 0,106 | 0,127 |
|  |  | hard > easy vWM | -0,02 | 0,04 | -0,59 | -0,11 | 0,559 | 0,706 |
|  | MD | intact > degraded | -0,07 | 0,05 | -1,54 | -0,28 | 0,126 | 0,347 |
|  |  | hard > easy vWM | 0,37 | 0,07 | 5,12 | 0,93 | 0,000 | 0,000* |
| RightIFGoper | Language | intact > degraded | 0,11 | 0,05 | 2,40 | 0,43 | 0,018 | 0,031* |
|  |  | hard > easy vWM | 0,05 | 0,04 | 1,11 | 0,20 | 0,268 | 0,535 |
|  | MD | intact > degraded | -0,01 | 0,04 | -0,17 | -0,03 | 0,866 | 0,866 |
|  |  | hard > easy vWM | 0,33 | 0,06 | 6,01 | 1,09 | 0,000 | 0,000* |
| RightIFGtr | Language | intact > degraded | 0,16 | 0,05 | 3,50 | 0,63 | 0,001 | 0,001* |
|  |  | hard > easy vWM | -0,02 | 0,04 | -0,54 | -0,10 | 0,589 | 0,706 |
|  | MD | intact > degraded | -0,03 | 0,06 | -0,58 | -0,11 | 0,561 | 0,824 |
|  |  | hard > easy vWM | 0,33 | 0,06 | 5,65 | 1,02 | 0,000 | 0,000* |
